# IL-2- and IL-4-dependent signaling play separate and sequential roles in the differentiation of GATA3^hi^ TH2 cells *in vivo*

**DOI:** 10.64898/2026.03.04.709484

**Authors:** Greta R. Webb, Caterina Carcó, Shiau-Choot Tang, Marie-Sophie Fabre, Sotaro Ochiai, Abbie Larson, Jodie Chandler, Kerry L. Hilligan, Evelyn Hyde, Sam I. Old, Olivier Lamiable, Franca Ronchese

**Author notes:** GRW current address: VIB Center for Inflammation Research, Ghent, Belgium. Correspondence: F Ronchese, Malaghan Institute of Medical Research PO Box 7060 Newtown, Wellington 6242 New Zealand.

## Abstract

Naïve CD4^+^ T cells differentiation into T-helper type 2 (TH2) cells requires Interleukin 2 (IL-2) and IL-4-dependent signaling as well as presentation of allergens by dendritic cells (DCs). The role of IL-2 and IL-4 in TH2 differentiation has been mostly studied *in vitro*, however, these models do not account for the heterogeneity of TH responses and bypass the DC-derived signals that are necessary *in vivo*. We used cytokine-blocking antibodies and IL-4Rα^KO^/IL-4Rα^WT^ mixed bone marrow chimeras to show that IL-4 signaling was not required for initial upregulation of GATA3 by CD4^+^ T cells after intradermal immunization, but was necessary for later TH2 cell expansion and further GATA3 upregulation. Single-cell transcriptomics and computational analyses confirmed that IL-4 signaling was not necessary for TH2 identity but promoted TH2 proliferation and expression of a pathogenic signature. Early IL-2 blockade prevented GATA3 upregulation without affecting TH2 proliferation or the differentiation of TH1 and TFH subsets after immunization. Adoptive transfer experiments showed that reduced competition for IL-4 *in vivo* drove extensive T cell proliferation and preferential expansion of the GATA3^hi^ population. Overall, our results suggest temporally and functionally distinct roles of IL-2 and IL-4 during TH2 differentiation: IL-2 is necessary for GATA3 upregulation, while IL-4 drives subsequent TH2 proliferation and licensing to effector activity. Therefore, IL-2 and IL-4 act in a co-ordinated manner to respectively promote TH2 differentiation, expansion, and effector commitment in the LN.

## Introduction

T-helper type 2 (TH2) cells are specialized effector CD4⁺ lymphocytes that protect against helminths but drive allergic disease. TH2 cells highly express the lineage regulator GATA3 and produce Interleukin 4 (IL-4), IL-5 and IL-13, which drive IgE, eosinophilia, mucus production, inflammation and tissue remodelling(1).

*In vitro*, IL-2 and IL-4 signaling are sufficient to induce TH2 differentiation and have been extensively studied(2). IL-4 acts as the canonical polarizing cytokine: it binds to the type-I IL-4 receptor composed of IL-4Rα and the common γ-chain (γc) and activates the transcription factor (TF) STAT6. STAT6 induces GATA3 expression (3–5) which, together with STAT5 and c-MAF, promotes IL-4 transcription (6,7). GATA3 also supports TH2 expansion, including via the TF Growth Factor Independent-1 (Gfi1)(8–10). IL-4–IL-4Rα signaling through STAT6 strengthens GATA3 and IL-4Rα expression, forming a positive feedback loop that stabilizes the TH2 phenotype (11). IL-2 complements the IL-4-STAT6 signaling pathway by cooperating with T cell receptor (TCR) signals to promote proliferation and establish TH2 competence through upregulation of the high-affinity IL-2 receptor (CD25/IL-2Rα–IL-2Rβ–γc)(12). Concurrently, IL-2–STAT5 signaling upregulates IL-4Rα, increasing T cell sensitivity to IL-4(13), and synergizes with GATA3 to induce IL-4 production (12).

While reductionist *in vitro* systems have been invaluable in dissecting the intricate cytokine and transcriptional network underlying TH2 differentiation, they cannot fully account for the complexity of *in vivo* immune responses. Firstly, TH2 cells, initially defined only by their ability to produce IL-4, were found to include phenotypically and functionally distinct subsets (14) that were unequally dependent on IL-4 for their *in vivo* differentiation (15) and effector activity (15–17). These include the IL-4^+^CXCR5^+^ type-2 T follicular helper cells (TFH2) cells which localize to B cell follicles (14) and help B cells switch to IgE production (18), and the GATA3^+^CCR4^+^ TH2 effectors expressing additional cytokines including IL-5 and IL-13 upon peripheral licensing (19). Secondly, *in vitro* models do not encompass signals from type-2 conventional dendritic cells (cDC2s) in shaping TH2 differentiation *in vivo* (20–26), the competition of CD4^+^ T cells for access to antigen-presenting cells (APCs) (27,28), or the cytokine production and consumption gradients that also control T cell activation and immune responses *in vivo* (29).

The key roles of IL-2 and IL-4 signaling in early TH2 responses in skin LN were recently examined using adoptive transfer of OVA-specific OT-II cells and intradermal immunization with OVA+papain. One study found that TH2 differentiation required paracrine IL-2- and IL-4-dependent signaling, which were supported by the LFA-1-dependent formation of DC-T cell “macro-clusters” at the T-B border area of the lymph node (30). Expression of pSTAT5 and pSTAT6 in antigen-specific CD4^+^ T cells in macro-clusters correlated with high GATA3 expression in the same cells (30). A second study found that CD301b^+^ cDC2s, which rapidly interact with incoming naïve CD4^+^ cells near high endothelial venules in LN (31,32), were an important source of the initial IL-2 for STAT5-dependent signaling in CD4^+^ T cells(33) to reduce BCL6 expression and promote differentiation into effector TH2 (33,34). Studies of intranasal exposure to the allergen House Dust Mite (HDM) also highlighted a key role of early IL-2 signaling within spatial “micro-niches” in mediastinal LN: IL-2 enabled GATA3 upregulation (35) and the TH2 effector program by suppressing BCL6 activity (34,36) through BLIMP-1 upregulation (37). IL-10 and STAT3-dependent signaling in HDM-specific CD4^+^ T cells also contributed to the permissive environment for BLIMP-1 upregulation by suppressing BACH2 expression (37).

While current evidence shows that IL-2 and IL-4 cooperate to promote effector TH2 differentiation while restraining TFH development, it is also known that these cytokines have complex effects as they both regulate their own and each other’s production and receptor expression (12,38–42). The temporal order of IL-2 and IL-4 expression within draining lymph node and their respective roles in directing CD4^+^ T cell differentiation into TH2 have not been fully dissected. Here we used antibody blockade, mixed bone-marrow chimeras, and adoptive transfer models to ask whether IL-2 and IL-4 control distinct checkpoints in TH2 differentiation *in vivo*. To capture the transcriptional dynamics of TH2 priming and expansion we performed single-cell (sc) RNA sequencing of lymph-node T cells during TH1 and TH2 immunization. Together, these approaches demonstrated a clear difference in the functional role of IL-2 and IL-4 signaling, with IL-2 necessary for early GATA3 upregulation, and IL-4 driving subsequent proliferation and functional specialization of TH2 effector cells. Thus IL-2 and IL-4 play distinct and complementary roles in the differentiation of CD4^+^ T cells into TH2 effectors *in vivo*.

## Results

### Expression of IL-4Rα and IL-2Rα are regulated by IL-4Rα-dependent signaling during early TH2 differentiation

To define the roles of IL-4 and IL-2 signaling in TH2 cell differentiation, we first characterized TH2 cells in skin-draining LN after intradermal immunization. By using a combination of intracellular and surface staining, we identified a distinct CD4^+^GATA3^high^ (hi) T cell population that was present in the LNs of mice immunized with non-viable *Nippostrongylus brasiliensis* L3 larvae (*Nb*) or HDM, but not after injection of PBS or heat-killed *Mycobacterium smegmatis* (*Ms*) (**Figures S1A, S2A-B**). This GATA3^hi^ population was partly IL-2Rα^hi^, expressed CCR4 but not CXCR5 (**Figure S2C**), and was not affected by the conditional deletion of BCL6 in T cells (**Figure S2D**) indicating that its development was BCL6-independent and therefore distinct from TFH.

To pinpoint the precise timing of IL-4 signaling on the differentiation of GATA3^hi^ cells, we immunized mice with *Nb* and treated with IL-4-neutralizing antibodies (aIL-4) or isotype control on day 2, 3 or 4 following immunization (**Figure 1A**). Anti-IL-4 treatment on day 2 did not affect the number of GATA3^hi^ cells on day 3 but significantly reduced them on day 5 (**Figure 1B**). Anti-IL-4 treatment on day 3 also reduced GATA3^hi^ numbers on day 5 but less effectively than aIL-4 treatment on day 2, while treatment on day 4 had no effect (**Figure 1B**). GATA3 levels also showed a trend to lower expression, but only on day 3 (**Figure 1C**). Anti-IL-4 treatment did not reduce the numbers of TFH cells compared to isotype, although a moderate but not statistically significant effect was observed when aIL-4 was given on day 2 for harvest on day 5 (**Figure 1D**).

**Figure 1:**
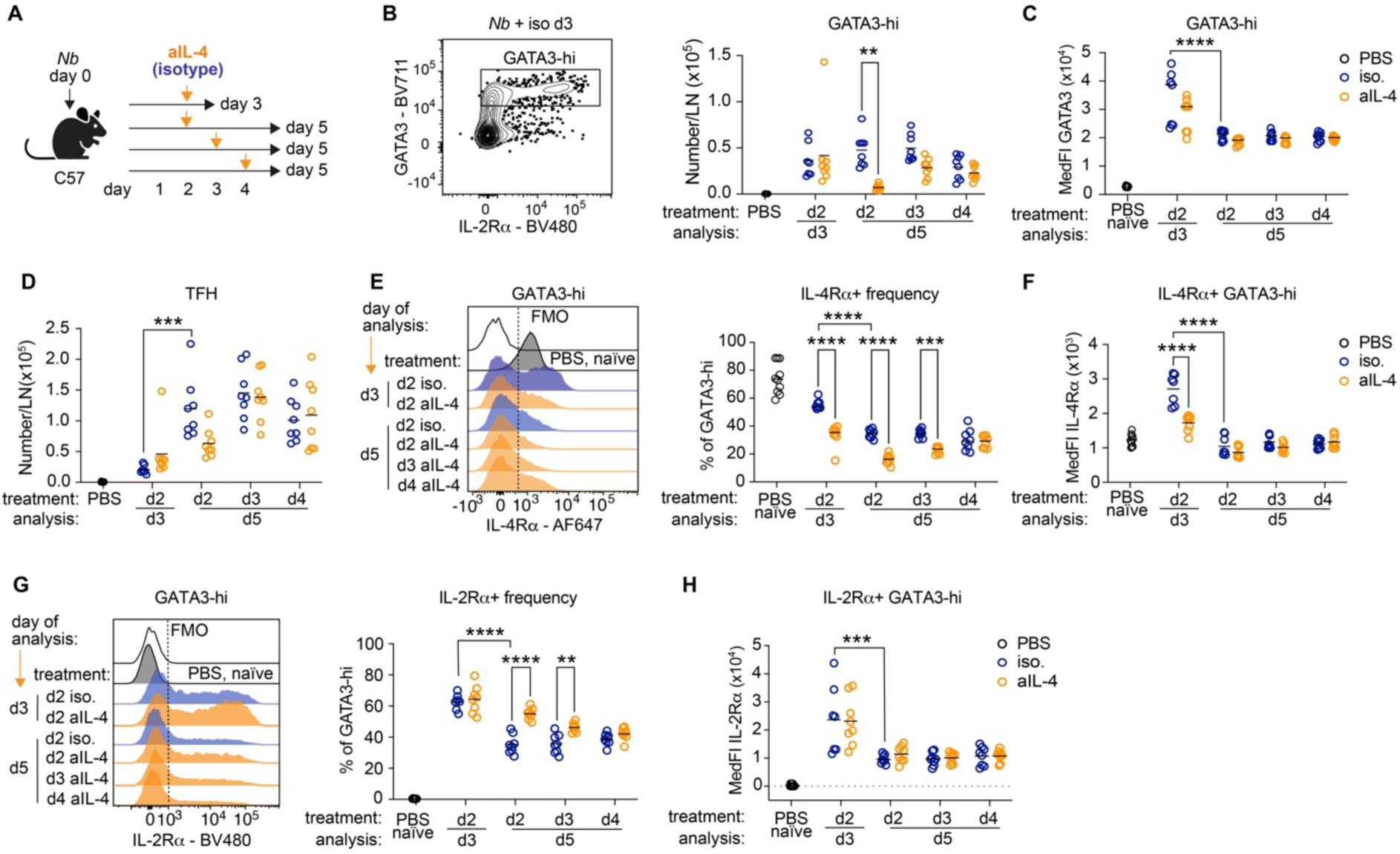
IL-4 blockade demonstrates a role for IL-4 signaling in the development of GATA3^hi^ CD4^+^ T cells following *Nb* immunization. C57BL/6 mice were immunized with *Nb* and treated with anti-IL-4 (aIL-4) or isotype control at the indicated times; CD4^+^ T cell responses in ear-draining LN were measured by flow cytometry on day 3 and 5 after immunization. **A:** Experimental schematic showing the timing of *Nb* immunization and aIL-4 treatment. **B:** Representative flow plot showing LN GATA3^hi^ CD4^+^ T cells following *Nb* immunization. Gating was as in Fig S1A. **C:** GATA3 Median fluorescence intensities (MedFIs) for the GATA3^hi^ T cell populations in (**B**). “PBS naïve” refers to PD1-low CD44-low FoxP3-CD4^+^cells from PBS-injected mice. **D:** Numbers of TFH cells per LN. **E:** Representative histograms and frequencies of IL-4Rα^+^ cells in GATA3^hi^ CD4^+^ T cell populations. The dotted line shows the gating for IL-4Rα^+^ cells. **F:** IL-4Rα MedFIs in the IL-4Rα^+^ T cell populations in (**E**). **G:** Representative histograms and frequencies of IL-2Rα^+^ cells in GATA3^hi^ CD4^+^ T populations. The dotted line shows the gating for IL-2Rα^+^ cells. **H:** IL-2Rα MedFI in the IL-2Rα^+^ T cell populations in (**G**). Data are pooled from two independent experiments, each with 3 – 5 mice per group, which gave similar results. Each symbol refers to one mouse. *P* values refer to the comparison between treatment groups and were calculated using two-way ANOVA with Tukey’s multiple comparisons test. ****, *P*<0.0001; ***, *P*<0.001; **, *P*<0.01. Only selected significant *P* values are shown.

We then examined IL-4Rα expression as it is upregulated in naïve CD4^+^ T cells upon bystander IL-4 exposure (40) but rapidly downregulated upon antigen recognition (39). IL-4Rα expression on GATA3^hi^ cells on day 3 was bimodal (**Figure 1E**), with about half of the GATA3^hi^ cells expressing high IL-4Rα whereas the remaining GATA3^hi^ cells were IL-4Rα low or almost negative. The proportion of IL-4Rα-hi cells declined to about 30% of GATA3^hi^ by day 5 (**Figure 1E**). Treatment with aIL-4-accelerated IL-4Rα downregulation: fewer GATA3^hi^ cells expressed IL-4Rα on day 3, and their level of expression was lower than in isotype-treated mice (**Figure 1E, F**). Therefore, IL-4-dependent signaling delays IL-4Rα downregulation on activated CD4^+^ T cells *in vivo*. In contrast to GATA3^hi^ cells, fewer than 20% of TFH expressed IL-4Rα on day 3, and fewer than 10% expressed it on day 5 (**Figure S3A**). This expression was further reduced, although not significantly, by aIL-4 treatment given on day 2. Anti-IL-4 treatment on day 2 and 3 also prevented IL-4Rα upregulation on CD44^low^PD1^neg^ CD4^+^ T cells, referred to as “naïve” hereafter, which otherwise expressed elevated IL-4Rα until day 5 (**Figure S3B**). Anti-IL-4 also reduced the modest IL-4Rα upregulation observed on Tregs (**Figure S3C**).

We then examined expression of IL-2Rα, as IL-2-dependent signaling is important for IL-4 production (2), GATA3 expression (43) and the acquisition of effector function by TH2 cells (34). Compared to naïve CD4^+^ T cells, IL-2Rα was highly expressed on GATA3^hi^ cells on day 3 and, although to a lower extent, on day 5 after *Nb* immunization (**Figure 1G**). Treatment with aIL-4 on day 2 and 3 after *Nb* immunization increased the frequency of IL-2Rα-expressing GATA3^hi^ cells on day 5 without affecting IL-2Rα MedFI (**Figure 1G, H**). This effect is consistent with the described inhibition of IL-2 production, and subsequent IL-2Rα expression, by IL-4 signaling (41,44).

These results suggest that IL-4-dependent signaling is required for the survival and/or expansion of already primed GATA3^hi^ TH2 cells in LN. IL-4 also maintains IL-4Rα expression on GATA3^hi^ cells while inhibiting IL-2Rα expression.

### STAT6 maintains IL-4Rα expression and suppresses IL-2Rα in GATA3-hi CD4+ T cells

As STAT6 is essential for IL-4 signaling, we examined TH2 responses and cytokine receptor expression in STAT6-deficient (STAT6^KO^) mice. GATA3^hi^ CD4^+^ T cells could be identified in STAT6^KO^ LN on day 3 after *Nb* immunization, albeit at lower numbers than in STAT6^WT^ (**Figure S3D**). GATA3^hi^ cell numbers on day 5 were significantly lower, and almost undetectable, in STAT6^KO^ compared to STAT6^WT^. T-bet^+^ T cell numbers following *Ms* immunization were similar on day 3 but lower in STAT6^KO^ than STAT6^WT^ on day 5 (**Figure S3E**), although this reduction was much less profound than the almost complete loss of GATA3^hi^ in STAT6^KO^ mice on day 5 after *Nb* (**Figure S3D**). TFH numbers following *Nb* or *Ms* were similar between STAT6^KO^ and STAT6^WT^ on both day 3 and day 5 (**Figure S3F**).

As also observed in **Figure 1E, F**, IL-4Rα expression on GATA3^hi^ T cells in STAT6^WT^ mice was rapidly upregulated following *Nb* immunization then gradually decreased (**Figure S3G, H**). By contrast, IL-4Rα expression on STAT6^KO^ GATA3^hi^ T cells was already low on day 3, and similar or lower than IL-4Rα expression on TFH (**Figure S3I**), or T-bet^+^CD4^+^ T cells in *Ms*-immunized STAT6^WT^ mice (**Figure S3G-I**), which rapidly downregulate IL-4Rα due to the lack of IL-4 production in this condition. We also examined IL-2Rα expression on GATA3^hi^ cells (**Figure S3J, K**): as observed in anti-IL-4 treated mice, the lack of IL-4 signaling in STAT6^KO^ GATA3^hi^ T cells prevented the downregulation of IL-2Rα, again suggesting an IL-4-dependent inhibition of IL-2 and IL-2-dependent IL-2Rα expression (41,44).

These results confirm the requirement for STAT6- and IL-4-dependent signaling in maintaining IL-4Rα expression on GATA3^hi^ T cells, and in downregulating IL-2Rα expression.

### IL-4Rα signaling is dispensable for the initial priming of GATA3^hi^ CD4^+^ T cells but is required for their expansion

To assess the T cell-intrinsic role of IL-4Rα signaling in TH2 differentiation, we generated mixed (IL-4Rα^WT^ + IL-4Rα^KO^) bone marrow (BM) chimeras which we immunized with *Nb* or PBS as a control (**Figure 2A**). Similar to the results with aIL-4 (**Figure 1B**), a distinct IL-4Rα^KO^ GATA3^hi^ population could be clearly identified in chimeric mice (**Figure 2B** and **S4A**). This GATA3^hi^ population expressed CCR4 (**Figure 2B**) and was moderately reduced compared to IL-4Rα^WT^ on day 3 (**Figure 2C**). On day 5, IL-4Rα^WT^ GATA3^hi^ cells had expanded in frequency compared to day 3, while IL-4Rα^KO^ CD4^+^ T cells had remained stable. IL-4Rα^WT^ GATA3^hi^ cells also expressed higher GATA3 compared to KO, although the increase was not statistically significant (**Figure 2C**), and comprised smaller proportions of IL-2Rα^+^ cells on both day 3 and day 5 (**Figure S4B**) to suggest reduced inhibition of IL-2 expression by IL-4 (42). In the spleen, IL-4Rα^WT^ GATA3^hi^ T cells outnumbered the IL-4Rα^KO^ (**Figure 2D**), excluding the possibility of differential egress of GATA3^hi^ cells from LN and entry into the circulation as an explanation for LN differences. Compared to IL-4Rα^WT^, IL-4Rα^KO^ TFH were slightly reduced in frequency on day 5 (**Figure 2E**), and they expressed lower levels of GATA3 compared to IL-4Rα^WT^ TFH (**Figure 2F**) thus reflecting the pattern observed in the GATA3^hi^ population.

**Figure 2:**
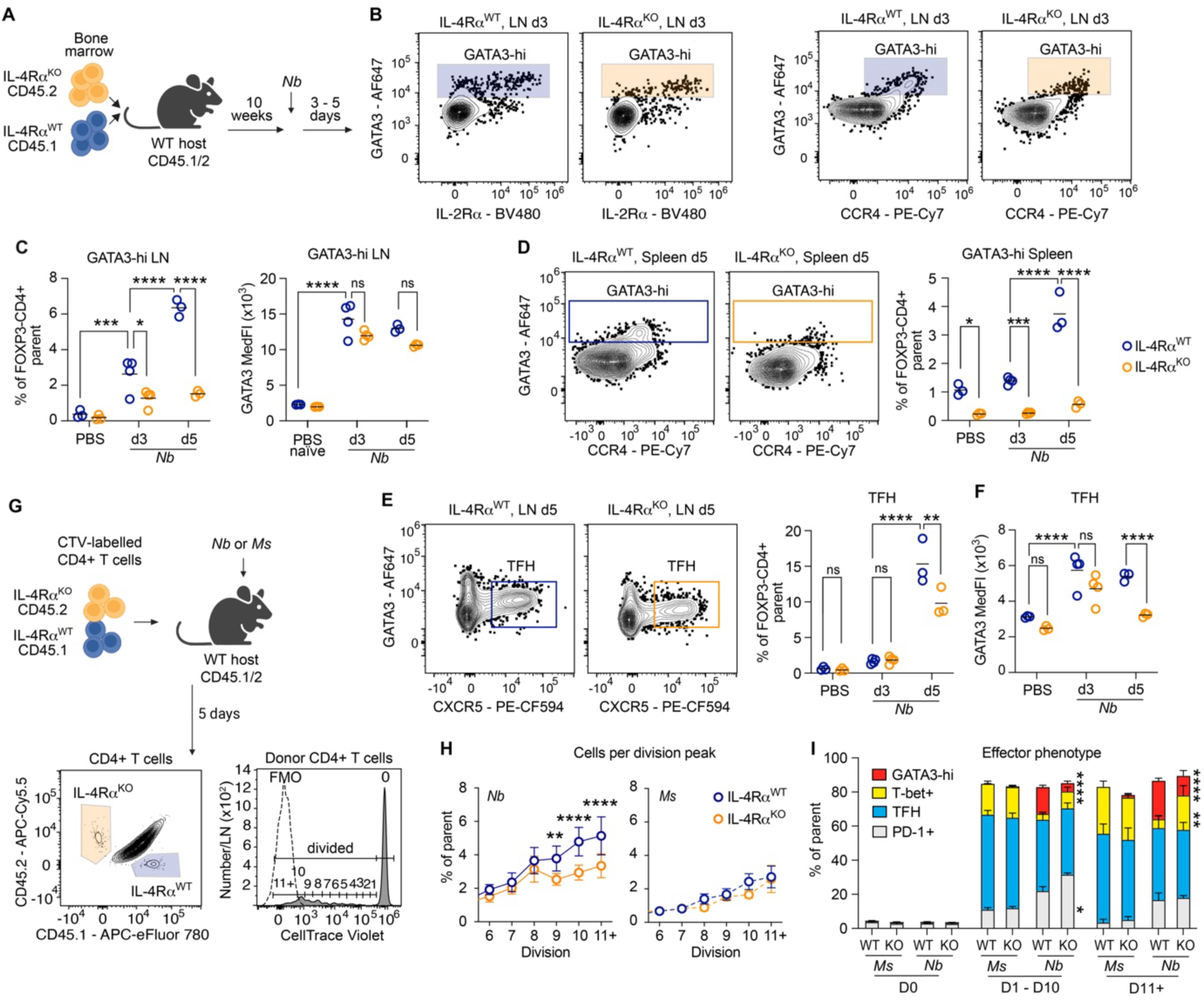
IL-4Rα signaling is required for GATA3^hi^ CD4^+^ T cell expansion. CD4^+^ T cell responses to *Nb* or *Ms* immunization in (IL-4Rα^WT^ + IL-4Rα^KO^) mixed BM chimeras (A-F), or C57xB6-SJ mice adoptively transferred with CellTrace Violet (CTV)-labelled IL-4Rα^WT^ or IL-4Rα^KO^ CD4^+^ T cells (G-I). **A:** Schematic showing the generation of mixed (IL-4Rα^WT^ + IL-4Rα^KO^) BM chimeras and timing of *Nb* immunization and analysis. **B:** Representative gating of GATA3^hi^ CD4^+^ T cells in ear-draining LNs as determined by surface and intranuclear staining. **C:** Frequencies of GATA3^hi^ CD4^+^ T cells in ear draining LN and their GATA3 median fluorescence intensities (MedFIs) in. “PBS naïve” refers to PD1-low CD44-low FoxP3-CD4^+^cells from PBS-injected mice. **D:** Representative gating and frequencies of GATA3^hi^ CD4^+^ T cells in spleen. **E:** Representative gating and frequencies of TFH in the LN. **F:** GATA3 MedFI in TFH from (E). **G:** Schematic and representative gating for *in vivo* proliferation assays. The histograms show division peaks of donor IL-4Rα^WT^ CD4^+^ T cells compared to CTV-unlabelled CD4^+^ T cells. **H:** Frequencies of donor CD4^+^ T cells per CTV division peak for *Nb* and *Ms*-immunized mice. Undivided cells and divisions 1 – 5 are not shown but were included in the frequency calculations. **I:** Phenotype of CD4^+^ T cells from undivided (0s), divided (1 – 10 divisions) and highly divided (11+ division) donor populations. Gating was as in Figure S1B, the remaining cells were PD-1-low and CD44-low and are not shown in the bar graph. All data are pooled from two independent experiments, each with 3 – 5 mice per group, which gave similar results. Symbols in **B-F** refer to individual mice; **H, I** show means and SEM. *P* values were calculated using two-way ANOVA with Tukey’s (B-F) or Šidák’s (H, I) multiple comparisons test. ****, *P*<0.0001; ***, *P*<0.001; **, *P*<0.01. Only selected significant *P* values are shown.

We then assessed cytokine expression by *in vitro* restimulation with PMA/ionomycin and intracellular staining. After *Nb* immunization, the frequencies of IL-4^+^CD4^+^ T cells in the IL-4Rα^WT^ and IL-4Rα^KO^ populations were similar on day 3 but lower in IL-4Rα^KO^ on day 5 (**Figure S4C)**. By contrast, the frequencies of IFNγ^+^CD4^+^ T cells following immunization with either *Nb* or *Ms* were similar between IL-4Rα^WT^ and IL-4Rα^KO^ on both day 3 and day 5 (**Figure S4D**). Therefore, IL-4Rα signaling is necessary for the development of IL-4^+^ TH2 and TFH cells but it is not required for IFNγ production.

To directly investigate whether IL-4 signaling has a role in CD4^+^ T cell expansion, we injected C57xB6.SJL host mice with a 1:1 mixture of IL-4Rα^WT^ and IL-4Rα^KO^ CD4^+^ T cells labelled with CellTrace Violet (CTV) (**Figure 2G**). While IL-4Rα expression was not necessary for CD4^+^ T cell division following *Nb* immunization, the proportion of cells that divided 9 or more times was higher in the IL-4Rα^WT^ population than in IL-4Rα^KO^ (**Figure 2H**). No difference in division was observed following immunization with *Ms* (**Figure 2H**). Further characterization of the dividing cells (**Figure S1B)** revealed that the majority of highly divided cells in *Nb*-immunized mice expressed T effector or TFH phenotypes, with a significantly higher proportion of GATA3^hi^ cells amongst the IL-4Rα^WT^ than IL-4Rα^KO^ donor cells at divisions 1-10 and 11+ (**Figure 2I**) but lower proportions of Tbet^+^ cells in division 11+ (**Figure 2I**). These data suggest that IL-4Rα signaling is not required for the differentiation of GATA3^hi^ CD4^+^ T cells but is essential for their proliferation and expansion.

### Effector CD4^+^ T cells from *Nb*- and *Ms*-immunized mice respectively express TH2 or TH1 transcriptional profiles

To obtain comprehensive information on IL-4Rα^KO^ CD4^+^ T cells compared to IL-4Rα^WT^ we carried out a scRNAseq analysis of activated (PD1^+^/CD44^+^) CD4^+^ T cells from mice immunized with *Nb* or *Ms* 3 days or 5 days earlier. Naïve (PD-1-low, CD44-low, CD62L^+^) cells from the same mice were collected as a comparison (**Figure S1C**). After quality control (QC), analysis of the integrated datasets revealed three broad populations (**Figure S5A, B**) that all expressed high levels of TCR genes (*Cd3d, Cd3e, Cd3g*), *Cd4* and *Lck*, but low *CD8a*, *CD8b1*, *Nkg7* and *Klrb1c* (**Figure S5C**), confirming their identity as CD4^+^ T cells. Clusters corresponding to Naïve (*Ccr7*, *Il4ra*, *Sell*), Activated (*Cd44*, *Pdcd1*), and Treg populations (*Foxp3*, *Il2ra*, *Ikzf2*) (**Figure S5C**) were also identified.

Re-clustering of the Activated compartment identified 13 clusters defined by marker expression and cell-cycle scores (**Figures 3A-C** and **S5D)**. We identified a cluster of non-cycling *Ccr7*^+^ *Bach2*^hi^ cells, and a small Recently-act(ivated) cluster characterized by high expression of the early activation markers *Cd69*, *Nr4a1* (NUR77), *Egr1-3*, *Irf4* (**Figure 3B**), and the TH0 cytokines *Il2*, *Il21* and *Tnf* (**Figure 3D**), whereas expression of the effector cytokines *Il4*, *Ifng* and *Il17a* was low or absent. Three clusters of cycling cells, Cycling-1, −2 and −3 expressed cell cycle genes (*Top2a*, *Mki67, Pclaf*, *Mettl1, Pus7 and Srm*) and high S and G2-phase scores. Two populations, TFH-1 and TFH-2 were identified as TFH cells. They both expressed *Cxcr5* and *Bcl6,* but the TFH-2 expressed lower *Bcl6* and *H2-Q2* together with the memory markers *Sell, S1pr1* and *S1pr4* (45), suggesting partial differentiation (46) or a transitional phenotype.

**Figure 3.**
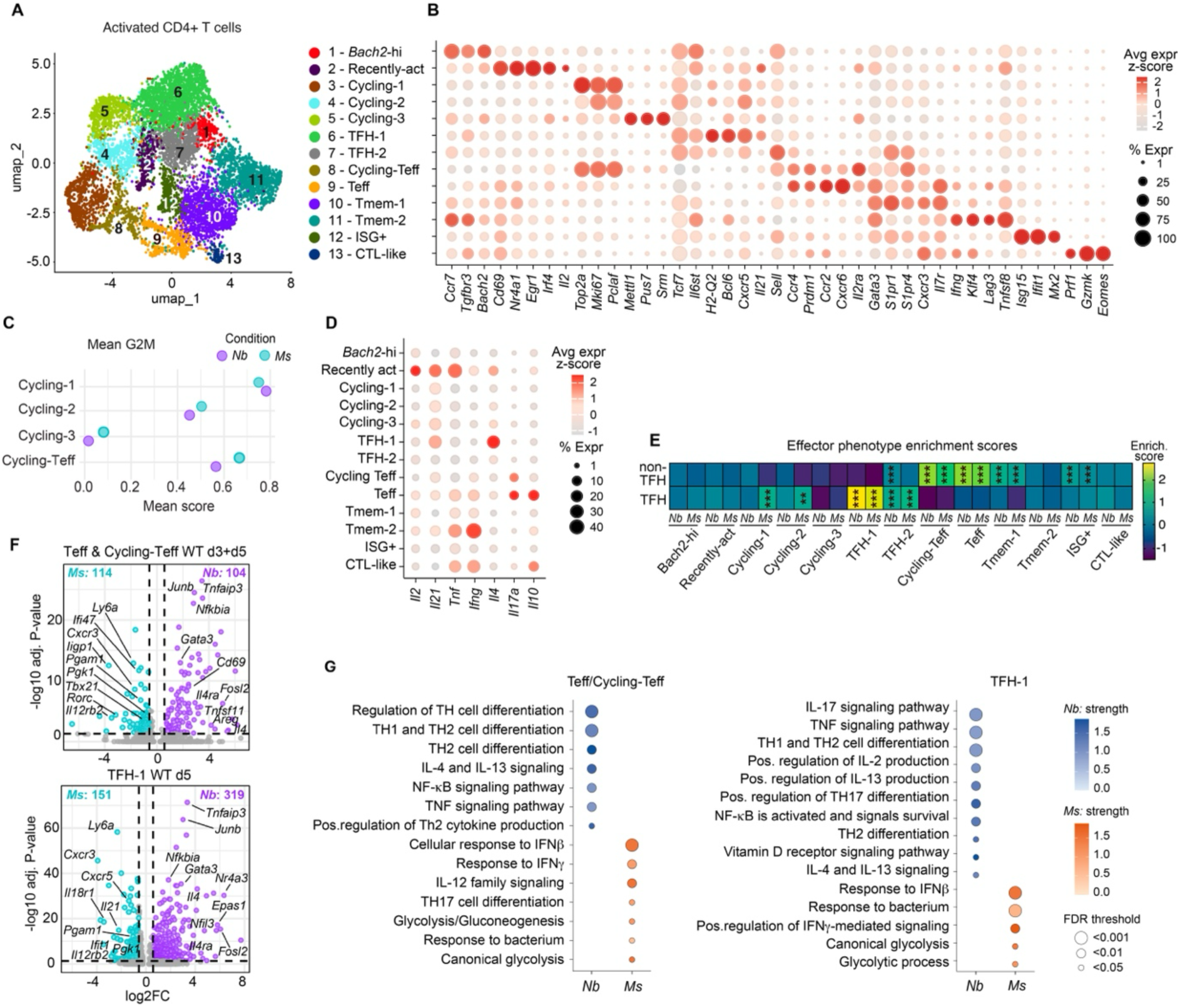
Transcriptomic characterization of activated CD4^+^ T cells from (IL-4Rα^WT^ + IL-4Rα^KO^) mixed BM chimeras following *Nb* or *Ms* immunization. Mixed-bone marrow chimeras were immunized with *Nb* or *Ms*. Naïve and activated IL-4Rα^WT^ and IL-4Rα^KO^ CD4^+^ T cells were FACS-sorted from ear draining LNs on day 3 and 5 following immunization according to the strategy in Figure S1B and processed for scRNAseq. **A:** Uniform manifold approximation and projection (UMAP) of CD4^+^ T cell clusters identified by scRNA-seq after QC filtering and exclusion of naïve and Treg cells as in Figure S5A. **B:** Average expression of cluster-defining genes for the 13 clusters in (**A**) in the total dataset (*Nb* and *Ms*, day 3 and day 5, WT and KO). Genes were curated from Seurat’s FindAllMarkers analysis (two-sided Wilcoxon rank sum test with Bonferroni correction) and were supplemented with relevant markers based on *a priori* knowledge. **C:** Mean G2M cell cycle phase scores for the indicated clusters and conditions; combined scores (day 3 and day 5, WT and KO) are shown. **D:** Average expression of selected cytokine transcripts for the 13 clusters in (**A**), in the total dataset. **E:** Enrichment of “non-TFH” or “TFH” gene signatures across the CD4⁺ T cell clusters in **A**. Significance was determined by comparing the score of each cluster to the score of all other clusters combined using Wilcoxon rank-sum tests with Benjamini–Hochberg correction. An FDR < 0.05 was considered significant. ***, FDR < 0.001; **: FDR < 0.01 are shown only for positive enrichment scores. **F:** Volcano plots comparing gene expression in IL-4Rα^WT^ Teff & Cycling-Teff and IL-4Rα^WT^ TFH-1 clusters from *Nb*- or *Ms*-immunized mice as indicated. **G**: Functional enrichment analysis of upregulated DEGs in the indicated IL-4Rα^WT^ clusters, comparing *Nb* and *Ms* conditions as in (**F**). Pathways were identified using the STRING database. FDR < 0.05 were considered significantly enriched. scRNAseq data are from one (*Ms*) or two (*Nb*) biological replicates.

Four non-TFH clusters, Cycling-Teffectors (Teff), Teff, Tmemory (Tmem)-1 and Tmem-2 variably expressed intermediate *Cd69,* and high *Gata3*, *Cxcr3* and *Il7r* (**Figure 3B**). The closely related Cycling-Teff and Teff were more abundant after *Nb* than *Ms* immunization on day 5 (**Figure S5E**) and uniquely expressed *Ccr4* and the *Bcl6* repressor *Prdm1*/BLIMP1, thus suggesting that this cluster comprised CD4^+^ T cells undergoing differentiation to effector function. They were distinguished from each other by the high expression of *Il2ra*, the cell-cycle genes *Top2a*, *Mki67* and *Pclaf*, and high G2M- and S-phase scores in the Cycling-Teff; and higher *Ccr2*, tissue residence marker *Cxcr6*, and cytokines *Il17a* and *Il10* in Teff (**Figure 3B, D**). Tmem-1 had a higher expression of *S1pr1* and *S1pr4*, as well as *Sell*, suggesting a central memory phenotype (45). Tmem-2 expressed *Ifng* and *Tnfsf8*/CD30L which are required for antibacterial immunity (47), as well as the TGFβ receptor family member *TgXr3* and the exhaustion marker *Lag3,* indicative of inhibitory signaling (48). We also identified an ISG^+^ cluster expressing high levels of IFN-stimulated genes (*Isg15*, *Ifit1* and *Mx2*), and a CTL-like cluster expressing *Prf1*, *Gzmk* and *Eomes*. Both phenotypes have been described in lymph nodes (49) and may be bystander populations, although ISG^+^ cells have been described as precursors to CD4^+^ central memory populations in other LN datasets (50).

To validate the assigned identities, we assessed expression of gene signatures generated from a bulk RNAseq dataset of *Il4*-AmCyan^+^ TFH and non-TFH cells from the LN of HDM-immunized *Il4*-AmCyan reporter mice (15,51). Cycling-Teff and Teff scored highly for non-TFH (**Figure 3E**). The Tmem-1 and ISG^+^ clusters also aligned with non-TFH, suggesting that they may include corresponding memory populations. TFH-1 from both *Nb*-immunized and *Ms*-immunized mice scored highly for the TFH signature, while *Nb* TFH-2 aligned with both signatures again suggesting a transitional phenotype (**Figure 3E**). All clusters were represented in both *Nb* and *Ms* datasets, but their proportions differed. Cycling clusters were more abundant in the *Ms* dataset on both day 3 and day 5, while *Bach2*-hi, Tmem1 and Tmem2 were enriched in the *Nb* dataset (**Figure S5E**).

We then compared gene expression between *Nb* and *Ms* conditions. *Bach2*-hi, Recently-act, TFH-2 and cycling clusters showed few upregulated DEGs, whereas the TFH-1, Tmem-1 and Tmem-2 clusters expressed many, especially on day 3 and more strongly following *Nb* than *Ms* immunization (**Figure S5F**), suggesting that the larger size of these clusters facilitated DEG detection.

To assess how immunization influenced DEG quality, the TFH-1 cluster which increased in frequency between day 3 and day 5, and the pooled Teff+Cycling-Teff clusters which expressed the effector phenotype of interest, were compared between *Ms* and *Nb* conditions (**Figure 3F**). Data were filtered for IL4-Ra^WT^ donor cells to exclude potential confounding effects due to the lack of IL-4Rα signaling. In the Teff+Cycling-Teff population, DEGs included the expected higher *Gata3*, *Junb*, *Il4ra* and *Il4* for *Nb*, and higher *Tbx21*, *Il12rb2* and Interferon signaling genes (*Cxcr3*, *Ly6a*, *Ifi47*) for *Ms*. Several of the same DEGs were also found when comparing TFH-1 in *Nb* and *Ms* conditions, also including higher *Il4* for *Nb* and higher *Il21* for *Ms*. Gene-enrichment analysis of these identified DEGs with STRING (**Figure 3G**) confirmed that Teff+Cycling-Teff and TFH expressed canonical pathways as *Nb* was enriched in IL-4 signaling, TH2 differentiation and NF-kB activation genes; whereas *Ms* was enriched in Glycolytic metabolism, Interferon and IL-12R signaling (**Figure 3G**).

Overall, our dataset documents the early development of effector T cells, TFH, and memory subsets during both TH1 and TH2 responses while identifying key genes in this process.

### IL-4Rα-dependent signaling activates an effector transcriptional program in GATA3^hi^ “Teff” cells

We compared gene expression in Activated IL-4Rα^WT^ and IL-4Rα^KO^ clusters following *Nb* immunization. Violin plots of MAGIC-imputed genes of interest in IL-4Rα^WT^ populations confirmed that *Gata3* was higher in all Activated clusters compared to Naïve, and was highest in the Cycling-Teff and Teff clusters followed by Tmem-1 (**Figure 4A**). Cycling-Teff expressed high *Prdm1* and *Il2ra*, identifying them as the GATA3^hi^, IL-2Rα^+^ TH2 cells found by flow cytometry. Cycling-Teff also expressed some *Il4,* which was higher than in Teff, while *Il2* was expressed in Teff but not Cycling-Teff. *Il2* was also expressed in Recently-act and Cycling-3 cells as well as in Tmem-1, Tmem-2 and ISG^+^. *Il2* expression without translation has been reported in memory cells of TRM phenotype (52), therefore the contribution of the Tmem clusters to the IL-2-producing pool in LN remains unclear. *Il4* was also expressed in Recently-act and Cycling clusters, but the highest expression was in TFH-1 as reported (14) whereas *Il4ra* was low in all clusters except Naïve. A comparable pattern of gene expression was observed in the IL-4Rα^KO^ clusters (**Figure S6A**), although lower *Gata3* especially in the cycling clusters was noted to suggest a slower upregulation of this marker when IL-4-dependent signaling was not available.

**Figure 4.**
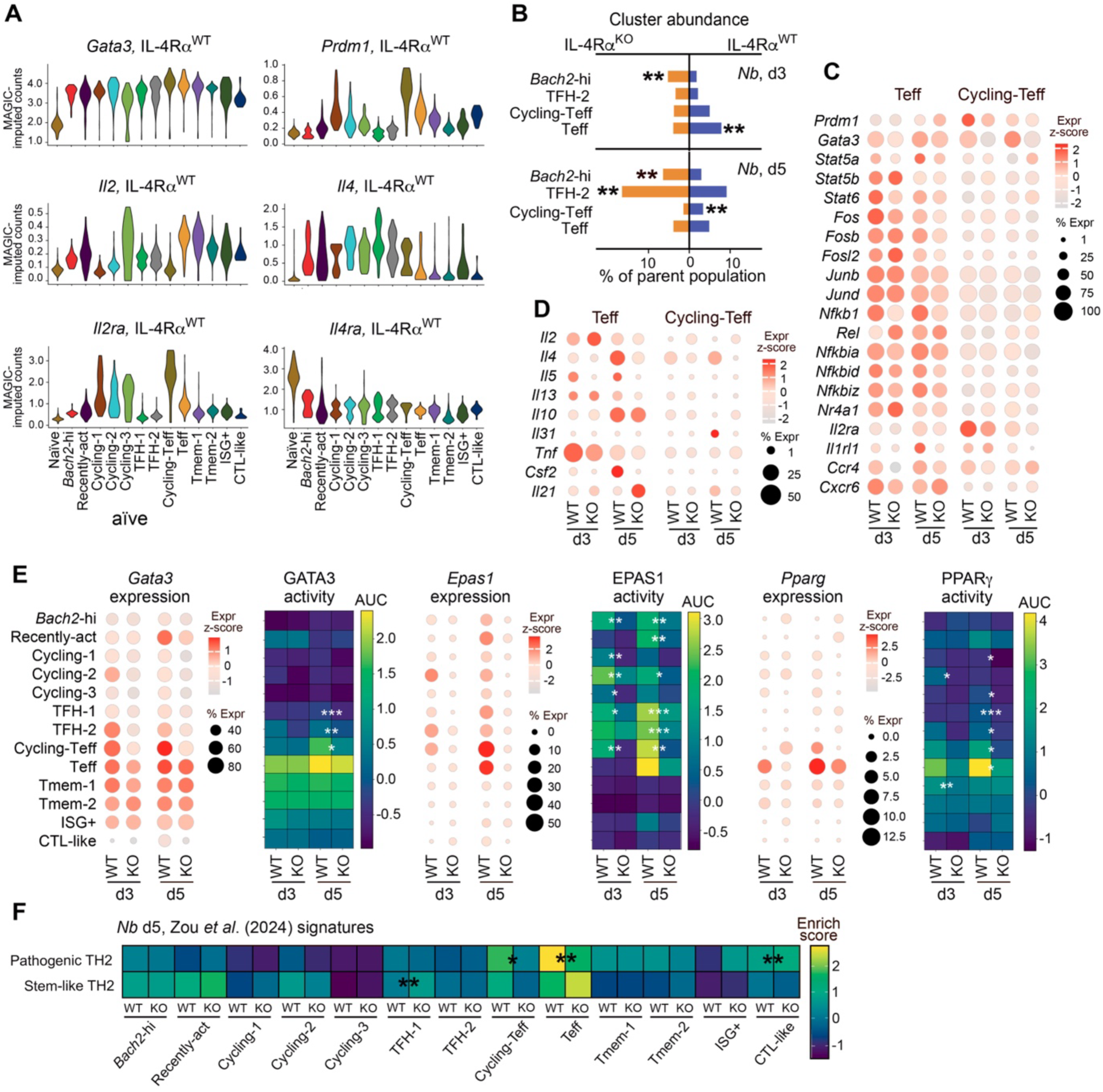
scRNA-seq analysis reveals expression of an IL-4Rα-dependent TH2 effector program in CD4⁺ T cells. (IL-4Rα^WT^ + IL-4Rα^KO^) mixed BM chimeras were immunized with *Nb*. Naïve and activated CD4^+^ T cells were FACS-sorted from the draining LN on day 3 and 5 following immunization according to the strategy in Figure S1B, and examined by scRNAseq. **A:** Violin plots show imputed single-cell expression levels of the indicated transcripts as generated with the MAGIC (Markov Affinity-based Graph Imputation of Cells) algorithm. Violins show combined day 3 and day 5 data in the IL-4Rα^WT^ population. **B:** Differences in selected cluster abundance between WT and KO. Significance was determined by permutation testing (the observed difference was compared with the difference expected by chance) and *P* values were adjusted for multiple testing using the Benjamini–Hochberg FDR procedure. Significant values were defined as log2FC > 0.58 and FDR < 0.05. **: FDR < 0.01. **C:** Expression of selected transcripts and **D**: cytokines in the Teff and Cycling-Teff clusters, by genotype and time following immunization with *Nb*. **E:** Expression and regulon activity (z-scored area under the curve, AUC) of TH2-associated transcription factors in IL-4Rα^WT^ and IL-4Rα^KO^ T cell clusters. **F:** Enrichment of “pathogenic” and “stem-like” TH2 gene signatures (from Zou *et al.,* 2024) in IL-4Rα^WT^ and IL-4Rα^KO^ T cell clusters. Differential regulon activity between WT and KO in **E**, **F** was assessed using a two-sided Mann–Whitney U test with Benjamini–Hochberg FDR correction for multiple comparisons. *: FDR < 0.05; **: FDR < 0.01; **\*\*\***: FDR < 0.001) are shown for positive enrichment scores. All data are pooled from two biological replicates.

We then examined how IL-4Rα expression influenced cluster representation in *Nb*-immunized chimeras. Only 4 clusters were significantly different between IL-4Rα^WT^ and IL-4Rα^KO^ on either day 3 or day 5. The *Bach2-*hi cluster was more abundant in the IL-4Rα^KO^ dataset compared to IL-4Rα^WT^ on both day 3 and day 5 following *Nb* immunization (**Figure 4B**), and on day 5 following *Ms* immunization (**Figure S6B**), suggesting that the developmental progression of the corresponding cells required IL-4Rα-dependent signaling. The heterogeneous TFH-2 cluster was also over-represented in the IL-4Rα^KO^ dataset on day 5 following *Nb* immunization (**Figure 4B**), again suggesting that IL-4Rα signaling promotes progression into mature phenotypes. In contrast to *Bach2*-hi and TFH2, Teff were more abundant in the IL-4Rα^WT^ dataset on day 3, possibly suggesting a delayed acquisition of effector phenotype by IL-4Rα^KO^ T cells (**Figure 4B**), and IL-4Rα^WT^ cells comprised the majority of the Cycling-Teff population on day 5, confirming that IL-4Rα signaling is required for TH2 effector expansion (**Figure 4B**). The IL-4Rα^WT^ and IL-4Rα^KO^ Teff clusters were similarly represented in *Ms*-immunized mice (**Figure S6B**), demonstrating that the reduced effector differentiation was specific to TH2 conditions. Finally, RNA velocity (**Figure S6C**) and pseudotime (**Figure S6D**) analyses revealed no major differences in trajectories between IL-4Rα^WT^ and IL-4Rα^KO^ cells after *Nb* immunization (p = 0.412 for pseudotime analysis in **Figure S6D**) suggesting that loss of IL-4Rα signaling alters the efficiency of effector acquisition rather than the overall TH2 developmental pathway.

We focussed on Teff and Cycling-Teff as they showed the highest expression of TH2-associated markers (**Figures 3B, E** and **4A**) and were less abundant in IL-4Rα^KO^ than IL-4Rα^WT^ (**Figure 4B**). We first examined the expression of selected *Nb*-associated DEGs within these clusters (**Figure 4C**). *Prdm1* was highly expressed in Cycling-Teff cells and showed reduced expression in IL-4Rα^KO^ cells especially on day 3. *Gata3* expression was also reduced in IL-4Rα^KO^ in both clusters and both days. These results confirm that, in the absence of IL-4Rα signaling, CD4^+^ T cells have a reduced capacity to develop into GATA3^hi^ effector cells. *Stat5a* and *Stat5b* were both expressed, with *Stat5b* more highly expressed on day 3 and *Stat5a* on day 5 and preferentially in WT, while *Stat6* was expressed early and only in WT as expected (**Figure 4C)**. Several AP-1 family transcription factors previously linked to pathogenic TH2 cells (53), *Fos*, *Fosb* and *Fosl2*, were also expressed with a trend to higher expression in IL-4Rα^WT^ Teff clusters, while no differences were observed for *Junb* and *Jund*. In addition, compared to IL-4Rα^WT^, IL-4Rα^KO^ Teff cells showed a marked reduction in expression of *N[b1*/p50, which was recently reported to be required for TH2 effector function in the lung (54). This reduction was accompanied by decreased expression of the NF-kB signaling inhibitors *N[bia, N[bid and N[biz* on day 5, likely reflecting impaired negative feedback regulation in the absence of IL-4Rα signaling (**Figure 4C**). Consistent with effector differentiation, the Teff compartment showed an enrichment of the IL-33 receptor *Il1rl1* and the chemokine receptor *Ccr4*, together with the cytokines *Il4*, *Il5*, and *Csf2* (**Figure 4D**), which were all preferentially associated with the IL-4Rα^WT^ population on day 5 (**Figure 4C-D**). Conversely, the IL-4Rα^KO^ Teff cluster showed increased expression of *Nr4a1*, a transcription factor known to reduce CD4^+^ T cell activation by restricting early metabolic reprogramming (55).

Our comparison of the transcriptional profiles of IL-4Rα^WT^ and IL-4Rα^KO^ cells suggested that IL-4Rα signaling regulates gene programs associated with TH2 effector function. To investigate this further, we performed gene regulatory network analysis on Teff and Cycling-Teff cells from IL-4Rα^WT^ and IL-4Rα^KO^ populations on day 3 and day 5, focusing on established transcriptional regulators of TH2 differentiation. Notably, the regulons for PPARG and EPAS1, two key regulators of the pathogenic TH2 program (56–58), showed high activity in both Teff and Cycling-Teff clusters on day 5, but their activity was restricted to the IL-4Rα^WT^ cells (**Figure S6E**). Consistent with this observation, the expression of *Epas1* and *Pparg* was highest in Teff and Cycling-Teff compared to other clusters and required IL-4Rα-dependent signaling (**Figure 4E**). Analysis of GATA3, EPAS1 and PPARG transcriptional activities across all clusters recapitulated these findings (**Figure 4E**), suggesting that reduced *Epas1*/*Pparg* expression and/or activity in IL-4Rα^KO^ cells may contribute to impaired maturation of TH2 effector subsets. Gene enrichment analysis further supported this interpretation. Using published “stem-like” and “pathogenic” TH2 gene signatures from Zou *et al*. (57) we found that Teff and Cycling-Teff displayed a stronger pathogenic signature in IL-4Rα^WT^ than IL-4Rα^KO^ on day 5, whereas TFH-1 were enriched for the stem-like signature in IL-4Rα^KO^ (**Figure 4F**). These results indicate that IL-4Rα-dependent signaling is necessary for the acquisition of pathogenic potential.

Together, these analyses suggest that IL-4Rα-dependent signaling in TH2 cells is not necessary for GATA3 expression and activity, but supports TH2 progression into effector cells by sustaining key transcriptional programs controlled by EPAS1 and PPARG.

### IL-4Rα-dependent competition for IL-4 access controls CD4^+^ T cell proliferation in LN

To obtain further evidence on the role of IL-4Rα signaling in TH2 differentiation and T cell expansion *in vivo*, we employed an adoptive transfer model (**Figure 5A**) in which CTV-labelled IL-4Rα^WT^ CD4^+^ T cells were injected i.v. into either IL-4Rα^WT^ hosts, in which the ability of donor and host cells to bind IL-4 was similar, or into IL-4Rα^fl/-^ hosts in which endogenous cells expressed half the WT levels of IL-4Rα due to one null allele (**Figure S7A**) and therefore had lower capacity for IL-4 signaling than donor cells. This prediction was confirmed *in vitro* by showing that IL-4Rα^fl/-^ CD4^+^ T cells exposed to the same concentration of IL-4 in culture were less able to phosphorylate STAT6 compared to IL-4Rα^WT^, while IL-4Rα^KO^ cells did not phosphorylate STAT6 at all (**Figure S7B**).

**Figure 5:**
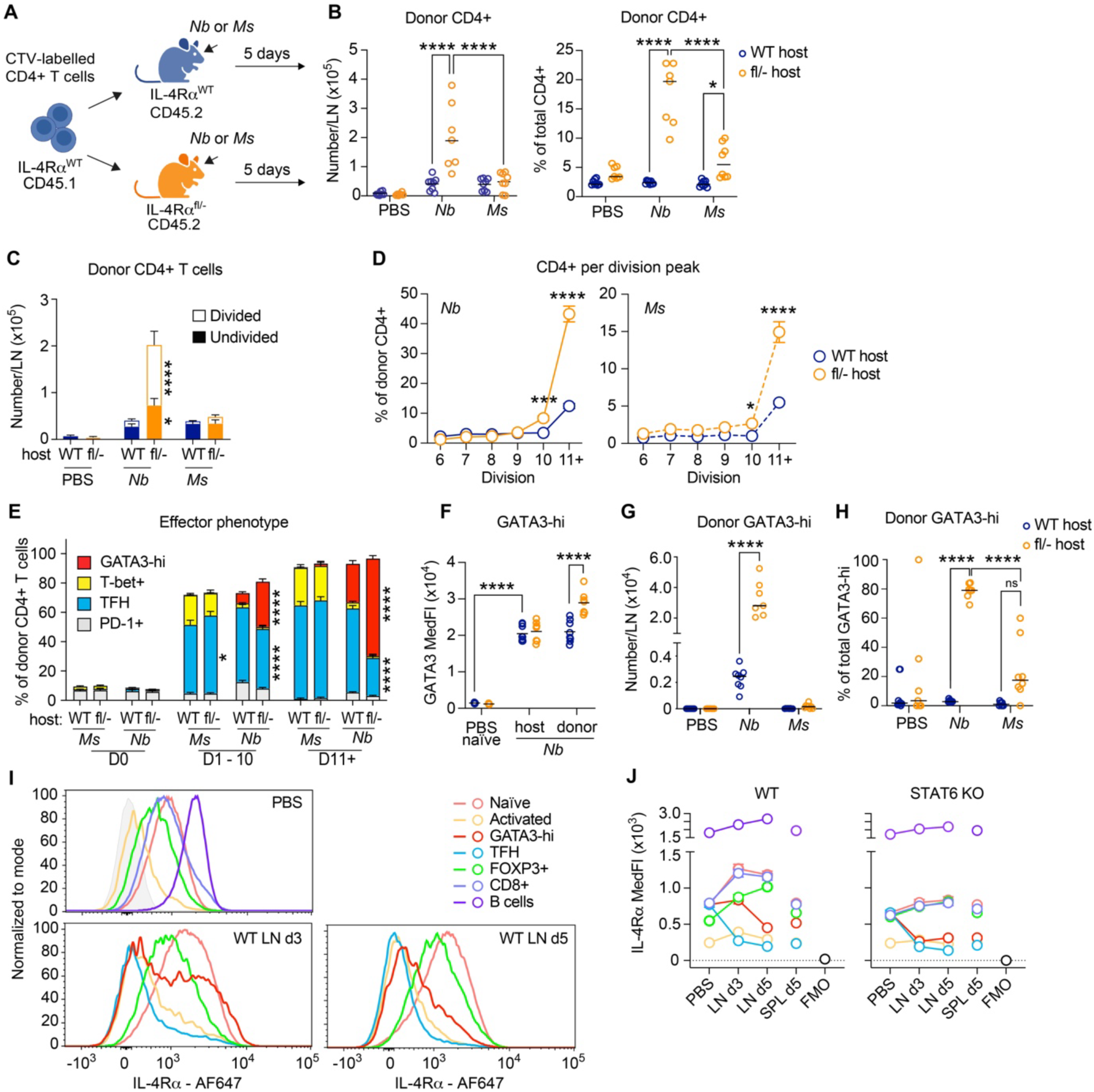
Reduced competition for IL-4 enables increased proliferation and preferential expansion of GATA3^hi^ CD4^+^ T cells. IL-4Rα^WT^ and IL-4Rα^fl/-^ hosts were adoptively transferred with CellTrace Violet (CTV)-labelled IL-4Rα^WT^ CD4^+^ T cells and immunized with *Nb* or *Ms*; CD4^+^ T cell responses in ear-draining LN were measured 5 days following immunization. **A:** Schematic for adoptive transfer and *in vivo* proliferation assay. **B:** Numbers and frequencies of donor CD4^+^ T cells per lymph node (LN) in the total CD4^+^ T cell population. **C:** Numbers of undivided and total divided donor CD4^+^ cells per LN. **D:** Frequencies of donor CD4^+^ T cells in each division peak following immunization with *Nb* or *Ms*. Undivided cells and divisions 1 – 5 are not shown but were included in the frequency calculations. **E:** Phenotype of effector CD4^+^ T cells in the undivided (D0), divided (D1 – 10) and highly divided (D11+) donor CD4^+^ T cell populations, gated as in Fig S1B. **F:** Median fluorescence intensity (MedFI) of GATA3 expression in GATA3^hi^ donor and host CD4^+^ T cells in adoptively transferred mice. **G:** Numbers of donor GATA3-hi CD4^+^ T cells per LN and (**I**) their frequencies within the indicated GATA3-hi populations. “PBS naïve” refers to CD44^low^PD-1^low^CD4^+^ T cells in PBS-treated mice. **H:** Flow cytometry histograms and (**K**) MedFIs of IL-4Rα expression in LN and spleen total populations. Fluorescence-minus-one (FMO) is shown in solid light grey only in the PBS panel. Symbols in **B, F - H** refer to individual mice; **C – E** and **J** show mean and SEM. All figures show pooled data from two repeat experiments, each with 3 – 5 mice per group, which gave similar results. *P* values were calculated using ordinary two-way ANOVA with Tukey’s or Šidak’s (**D** only) multiple comparisons test. Selected significant comparisons are shown. ****, *P*<0.0001; ***, *P*<0.001; **, *P*<0.01; *, *P*<0.05; ns, *P*≥0.05.

Adoptively transferred mice immunized with *Nb* or *Ms* were all able to generate GATA3^hi^, T-bet^+^ and TFH responses to both *Nb* and *Ms* immunization (**Figure S7C-E**), although GATA3^hi^ and TFH responses to *Nb* were significantly higher in WT hosts than in IL-4Rα^fl/-^ thus suggesting a role for IL-4 signaling in regulating the magnitude of these responses. The contribution of donor CD4^+^ T cells to these responses was host dependent: donor cells were recovered in higher numbers and proportions from *Nb*-immunized IL-4Rα^fl/-^ than IL-4Rα^WT^ hosts (**Figure 5B**) due to greater numbers of divided CD4^+^ T cells (**Figure 5C**) and higher frequencies of highly divided cells in both *Nb*- and *Ms*-immunized Il4ra^fl/-^ hosts compared to IL-4Rα^WT^ (**Figure 5D**). Further characterization of divided donor cells revealed a marked preferential expansion of GATA3^hi^ CD4^+^ T cells compared to all other activated subsets in *Nb*-immunized IL-4Rα^fl/-^ hosts, which was especially prominent in the highly divided (D11+) population (**Figure 5E**). Increased IL-4Rα signaling in GATA3^hi^ CD4^+^ T cells also led to increased GATA3 expression in GATA3^hi^ donor cells compared to host cells in IL-4Rα^fl/-^ mice (**Figure 5F**). A comparison of donor cell numbers and their proportions within each effector population showed that GATA3^hi^ cells underwent an about 20-fold greater expansion in IL-4Rα^fl/-^ hosts compared to IL-4Rα^WT^ (**Figure 5G**) to represent almost 80% of the GATA3^hi^ population in IL-4Rα^fl/-^ hosts. T-bet^+^, TFH, (**Figure S7F, G**) as well as IL-4, IFNγ and IL-2-expressing cells (**Figure S7H – M**) in *Nb*- and *Ms*-immunized mice were also preferentially expanded in IL-4Rα^fl/-^ hosts.

These results indicate that the ability to successfully compete for available IL-4, and signal through IL-4Rα, is a key mechanism that controls TH2 proliferation and GATA3 expression *in vivo*. To identify which populations may be competing against TH2 for IL-4 access, we examined IL-4Rα expression in LN. Following *Nb* immunization, IL-4Rα was progressively downregulated in both TH2 and TFH, whereas its average expression on naïve CD4^+^ T cells, CD8^+^ T cells, B cells and also FOXP3^+^ Tregs was upregulated in a STAT6-dependent fashion (**Figure 5I, J**), becoming at least two-fold higher than on TH2 cells on day 5. This two-fold competitor advantage would be lost in IL-4Rα^fl/-^ hosts, in which all bystander cells expressed levels of IL-4Rα that are half of WT. Although any of the IL-4Rα^hi^ populations could plausibly be competing for IL-4 with TH2, Tregs appear a relevant candidate given their reported proximity to IL-2^+^ CD4^+^ T cells undergoing activation in LN (59,60) and ability to compete for IL-2 via high IL-2Rα expression (61).

Together these results reinforce the key role of IL-4 in the expansion of GATA3^hi^ cells and GATA3 upregulation, and demonstrate a bystander IL-4Rα-dependent mechanism controlling IL-4 bioavailability and function *in vivo*.

### Signaling through IL-2Rα is critical for the emergence of GATA3^hi^ CD4^+^ T cells

As loss of IL-4Rα signaling reduced but did not prevent the differentiation of GATA3^hi^ CD4^+^ T cells, we explored the role of IL-2 signaling by treating mice with an anti-IL-2 neutralizing antibody given i.p. immediately after immunization with *Nb* or *Ms*. Compared to isotype control, anti-IL-2 treatment led to a striking reduction in GATA3^hi^ CD4^+^ T cells on both day 3 and 5 following *Nb* immunization (**Figures 6A, B** and **S8A**) with no significant effect on T-bet, TFH (**Figure 6C, D**) and Treg (**Figure S8B**) numbers. As expected, the frequencies of IL-2Rα^+^ cells in the GATA3^hi^ population and their IL-2Rα MedFI were both significantly reduced after aIL-2 treatment (**Figure 6E**). IL-4Rα expression on the few remaining GATA3^hi^ cells was also reduced (**Figure 6F**) although naïve CD4^+^ cells in the same LN upregulated IL-4Rα expression regardless of aIL-2 treatment (**Figure 6G**), thus indicating that IL-4 was being produced in the LNs of both control and anti-IL-2 treated mice (40).

**Figure 6:**
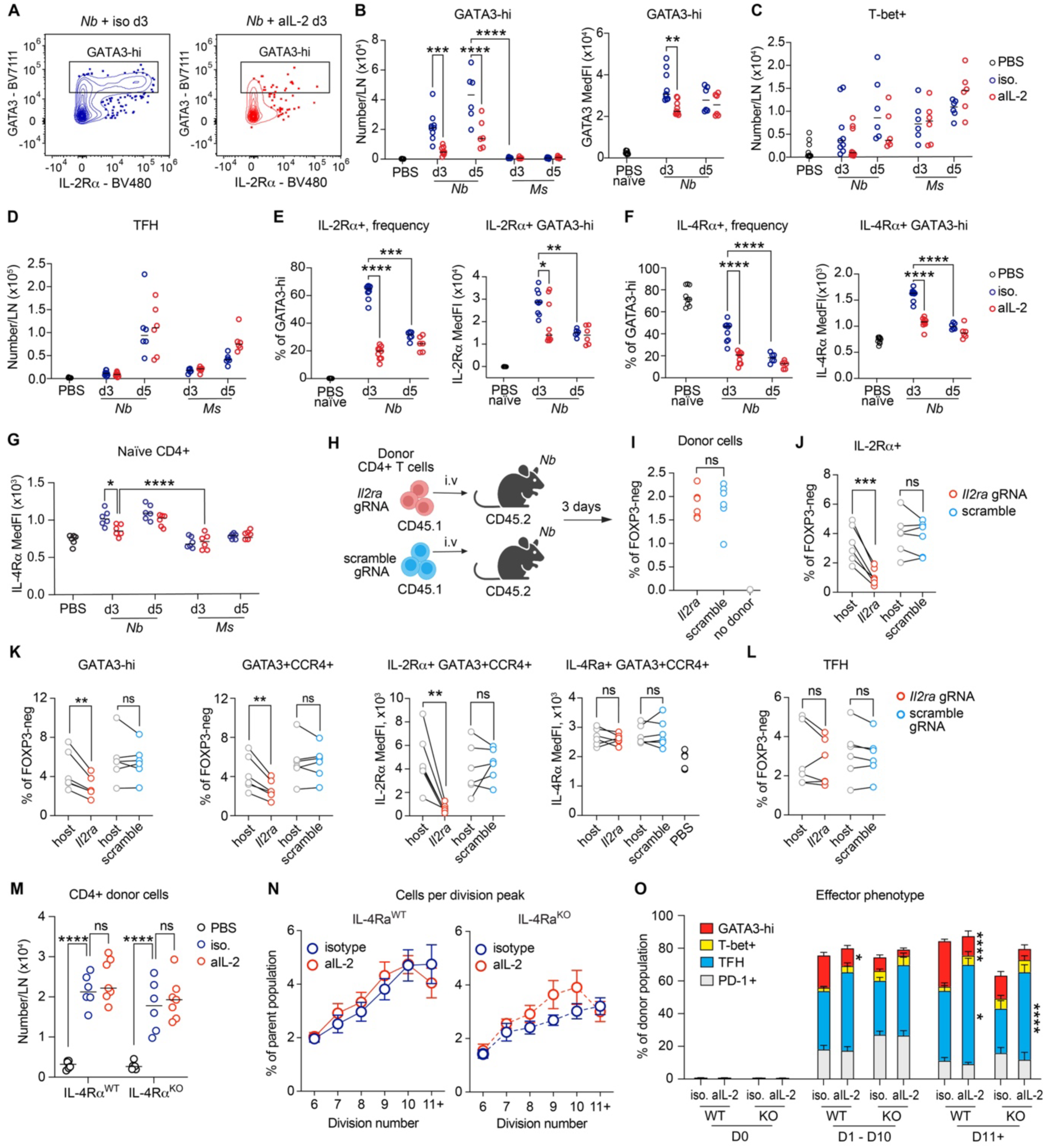
IL-2 signaling is essential for GATA3 upregulation and is not required for CD4^+^ T cell proliferation. C57BL/6 mice were immunized with *Nb* or *Ms*; CD4^+^ T cell responses in ear-draining LN were measured by flow cytometry at the indicated times after immunization. **A:** Representative gating of GATA3^hi^ CD4^+^ T cells. **B:** Numbers per lymph node (LN) and (**C**) GATA3 median fluorescence intensity (MedFI) of GATA3^hi^ CD4^+^ T cells. **C:** Numbers of T-bet^+^CD4^+^ T cells per LN. T-bet gating was as in Supplemental Figure 1A. **D:** Numbers of TFH cells per LN. **E:** Frequencies of IL-2Rα^+^ cells in the GATA3^hi^ CD4^+^ T cell population, and their IL-2Rα MedFI. IL-2Rα^+^ gating was as in Figure 1G. “PBS naïve” refers to CD44^low^PD-1^low^CD4^+^ T cells in PBS-treated mice. **F:** Frequencies of IL-4Rα^+^ cells in the GATA3^hi^ CD4^+^ T cell population and their IL-4Rα MedFI. IL-4Rα gating was as in Figure 1E. **G**: IL-4Rα MedFI in naïve CD4^+^ T cells from C57BL/6 mice immunized with *Nb* and treated with aIL-2 or isotype control. **H:** Experimental schematic for CRISPR knockdown experiments. **I**: Frequencies of donor cells in the CD4^+^FoxP3^neg^ (non-Treg) CD4^+^ T cell population. **J:** Frequencies of IL-2Rα^+^ T cells within host and donor CD4^+^FoxP3^neg^ populations. **K:** Frequencies of GATA3^hi^ and GATA3^+^CCR4^+^ TH2 cells, and their IL-2Rα and IL-4Rα expression. Gating was as in Figure 1E, G **L:** Frequencies of TFH cells within host and donor CD4^+^FoxP3^neg^ populations. **M:** Numbers of CD4^+^ of donor cells and (**N**) their frequencies by CTV division peak in mice immunized with *Nb* and treated with anti-IL-2. Undivided cells and divisions 1 - 5 are not shown in J but were included in the frequency calculations. **O:** Frequencies of effector CD4^+^ T cell subsets in divided donor CD4^+^ T cell populations. Experimental design and division gating were as in Figure 2G. Symbols in **B-G** and **I-M** refer to individual mice. Data are pooled from 2 or 3 repeat experiments, each with 3 - 6 mice per group, which gave similar results. Symbols and bar graphs in **N, O** show mean and SEM for two pooled experiments, each with 3 mice/group, that gave similar results. *P* values were calculated using two-way (**B-G, M-N**) or one-way (**O**) ANOVA with Tukey’s multiple comparisons test, or paired *t*-tests (**I-L**). ****, *P*<0.0001; ***, *P*<0.001; **, *P*<0.01. Selected *P* values are shown.

To directly assess the impact of anti-IL-2 on *Il4* expression we employed *Il4-*AmCyan (*Il4*-AmCy) reporter mice (62). Control and aIL-2-treated mice both generated *Il4*-AmCy^+^ CD4^+^ T cells following *Nb* immunization (**Figure S8C**), however, numbers in aIL-2-treated mice increased about one day later than in the isotype-treated controls, with both non-TFH and TFH *Il4*-AmCy^+^ CD4^+^ T cells comparably delayed (**Figure S8D-F**). Together, these data suggest that IL-4 was made in LN in both aIL-2 and isotype-treated mice, but its expression was delayed by aIL-2 treatment. Thus the accelerated decline of IL-4Rα expression on GATA3^hi^ and non-TFH cells was likely due to indirect effects of IL-2 blockade on IL-4 secretion, although direct IL-2 effects on IL-4Rα expression (13) were also possible.

To establish whether the effects of aIL-2 on GATA3^hi^ differentiation were due to cell-intrinsic effects on CD4^+^ T cells, we used CRISPR-mediated knockdown of *Il2ra* to disable high-affinity IL-2-dependent signaling in naïve CD4^+^ T cells, then adoptively transferred them into C57BL/6 hosts before *Nb* immunization (**Figure 6H**). CD4^+^ T cells targeted with scramble RNA were used as controls. Both CD4^+^ T cell populations survived similarly after transfer *in vivo* (**Figure S1D** and **6I**), and the *Il2ra* targeted cells expressed lower IL-2Rα than the host population (**Figure 6J**) indicating successful deletion. The *Il2ra*-targeted population generated lower frequencies of GATA3^hi^ and GATA3^+^CCR4^+^ cells (gated as in **Figure S2C**) which expressed lower IL-2Rα but similar IL-4Rα compared to the host population (**Figure 6K**), while the frequencies of TFH were not affected (**Figure 6L**). CD4^+^ T cells targeted with scramble RNA generated responses that were similar to the host population in all cases (**Figure 6J-L**). Therefore, IL-2Rα expression is intrinsically necessary for CD4^+^ T cell differentiation into GATA3^hi^, independently of any effects of IL-2Rα on Tregs, or on IL-4 production.

Finally, we investigated the role of IL-2 and IL-4 signaling in CD4^+^ T cell division using mice that were adoptively co-transferred with CTV-labelled IL-4Rα^WT^ and IL-4Rα^KO^ donor cells, immunized with *Nb,* and immediately treated with anti-IL-2 or isotype control. Treatment with aIL-2 did not significantly change the total number of adoptively transferred cells per LN (**Figure 6M**), or the frequency of cells at each division peak in either the IL-4Rα^WT^ or IL-4Rα^KO^ population (**Figure 6N**). Further characterization of dividing cells revealed that anti-IL-2 treatment led to a significant reduction in the frequency of GATA3^hi^ CD4^+^ T cells in the IL-4Rα^WT^ population, while TFH were increased in both D11+ IL-4Rα^WT^ and IL-4Rα^KO^ (**Figure 6O**). Therefore, IL-2 signaling is necessary for GATA3^hi^ T cell differentiation but does not affect T cell proliferation.

## Discussion

Here we show that IL-2 and IL-4 mediate functionally distinct roles in the differentiation of GATA3^hi^ TH2 effector cells in skin-dLN. Consistent with its early production in lymph node (33), IL-2 acted early and was essential for the upregulation of GATA3 but had limited or no role in the proliferation and expansion of TH2 cells. By contrast, IL-4-dependent signaling was not necessary for initial GATA3 expression but was a major driver of activated T cell division and expansion. While IL-4 could increase the expansion of multiple T cell populations in lymph node including Tbet^+^ CD4^+^ T cells and TFHs, its greatest impact was on GATA3^hi^ cells, which were progressively enriched in the proliferating CD4^+^ population and further upregulated GATA3 expression. In addition to increasing the number of GATA3^hi^ cells in LN, IL-4 also supported their expression of transcripts associated with TH2 effector function including low levels of the cytokines *Il5* and *Csf2*; the TH2-associated IL-33 receptor *Il1rl1* (ST2); and the TFs *Pparg* and *Epas1*. *Epas1*/HIF-2a was recently reported to be necessary for the transition from TH2 memory/stem to TH2 effectors (57), whereas PPARG sustains pathogenicity and the metabolic requirements of TH2 effectors in inflamed tissues (56,58). Both markers were predominantly expressed in TH2 effectors and were dependent on IL-4-dependent signaling, suggesting that IL-4 signaling in skin-dLN is already programming TH2 cells to effector function in the periphery.

Our studies differ from previous ones in several respects. Firstly, our experiments were carried out on polyclonal CD4^+^ T cell populations in WT C57BL/6 or IL-4Rα^KO^ donors. While this experimental design is the one that best reflects the heterogeneity and kinetics of a TH2 response in physiological settings, it also has some limitations. Firstly, the antigen specificity of our GATA3^hi^ CD4^+^ T cells was not formally proven. However, adoptive transfer of CTV-labelled polyclonal CD4^+^ T cells showed that high GATA3 expression was restricted to T cells that had divided multiple times to imply antigen-specific expansion. Secondly, our experimental design precluded the direct assessment of the earliest stages of TH2 differentiation on day 1 and 2 after immunization, when responding CD4^+^ T cells in dLN were too few for flow cytometric assessment. TH2 cells were clearly identifiable in the skin-dLN of *Nb*-immunized mice by day 3, when they had already differentiated into CCR4^+^IL-2Rα^+^GATA3^hi^, and by day 5 when they had also spread systemically as CCR4^+^GATA3^hi^ cells in spleen. Therefore, our work complements early activation experiments relying on the adoptive transfer of large numbers of TCR transgenic T cells of identical specificity (30,33,37) and extends those findings to a physiological setting and to include later phases of the immune response and development of effector phenotype. Secondly, many studies that assessed TH2 responses *in vitro* and *in vivo* relied on *Il4* transcriptional reporters as readouts. However, in LN *Il4* is expressed by both TH2 effectors and TFH (14,15), two populations with different functions, phenotypes and developmental requirements. We chose to focus specifically on TH2 effectors as mediators of allergic inflammation in tissues, and identified them through their high GATA3 expression as, together with IL-4, GATA3 is a defining markers of the TH2 lineage, necessary for sustained IL-4 production and TH2 responses (6,63,64). We confirmed that GATA3^hi^ cells were found in mice immunized with *Nb* or HDM but not after *Ms*; expressed the TH2 chemokine receptor CCR4 (65); and were also found in spleen to indicate the ability to spread systemically. GATA3^hi^ cells also lacked the TFH markers CXCR5, *Bcl6* and *Il21* and were found in similar numbers in BCL6^WT^ and BCL6^KO^ conditional KO mice (51) thus confirming their unique identity and developmental trajectory distinct from TFH.

We found that IL-4Rα expression was not necessary for the differentiation of GATA3^hi^ cells in LN, or for their expression of CCR4 which was also reported to require IL-4 for high expression and functionality (66). Although the levels of GATA3 were not as high in IL-4Rα^KO^ as they were in IL-4Rα^WT^ cells, they remained substantially higher than in other activated populations within the same LN, including TFH and Tregs. Accordingly, single-cell transcriptomic analyses indicated similar developmental trajectories for IL-4Rα^WT^ and IL-4Rα^KO^ populations, suggesting that factors other than IL-4 were driving high GATA3 expression and the development of TH2 identity in the IL-4Rα^KO^ population. While GATA3 upregulation did not require IL-4 signaling, IL-2 neutralization during *Nb* immunization, or IL-2Rα deletion in CD4^+^ T cells by CRISPR-Cas9 inactivation, led to a marked reduction in GATA3 and CCR4 expression following *Nb* immunization. IL-2Rα and STAT5-dependent IL-2 signaling play a key role in steering early CD4^+^ T cell fate (67) by controlling the reciprocal antagonism of the TFs BCL6, which is suppressed by IL-2 and favours TFH specification (68,69), and BLIMP-1 (encoded by *Prdm1*) which is IL-2 and IL-10 dependent (37) and necessary for effector activity and GATA3 upregulation in TH2 (35,70). Accordingly, in our dataset, high *Il2ra* expression in Cycling-Teff was associated with high *Prdm1* and low *Bcl6* and Bach2. While IL-2 signaling is clearly necessary for GATA3 expression and TH2 effector differentiation, as also shown by previous reports (34,37), it is important to note that additional signals are likely required. TCR and NOTCH-dependent signaling in CD4^+^ T cells have both been shown to upregulate GATA3 expression (reviewed in (71)). In addition, GATA3 expression in the absence of STAT6-dependent signaling was sufficient to auto-activate further GATA3 upregulation and induce TH2-specific chromatin remodelling (72). Any of these mechanisms may contribute to the GATA3 upregulation observed in our experiments.

*In vitro*, reduced GATA3 expression after IL-2 neutralization was hypothesized to be indirect and due to reduced IL-4 production(43) and/or IL-4Rα expression (13). Our *in vivo* experiments partly confirmed these findings by showing that IL-2 blockade at the time of immunization delayed IL-4-reporter expression *in vivo* by ∼24h. As expected on the basis of previous work (39), delayed IL-4 expression resulted in accelerated IL-4Rα downregulation during CD4^+^ T cell activation. By contrast, IL-2 signaling itself did not appear to be necessary for IL-4Rα expression, as CRISPR-Cas9 deletion of IL-2Rα in CD4^+^ T cells did not lead to reduced IL-4Rα expression upon adoptive transfer into an IL-4-sufficient environment. Together, these results suggest that lower expression of IL-4 and IL-4Rα are not sufficient to explain the lack of GATA3 upregulation following IL-2 blockade.

IL-4 is considered a key driver of TH2 differentiation and is necessary for IL-4 production *in vitro* (2) and TH2 response *in vivo* (16). Our data show that IL-4Rα deletion did not alter the core differentiation trajectory of TH2 cells and therefore are not consistent with a key role of IL-4 in initiating GATA3 upregulation and TH2 differentiation. However, our data do suggest that IL-4 has additional distinct but essential functions: IL-4 signaling was essential for vigorous proliferation and clonal expansion of GATA3^hi^ cells after activation, for further GATA3 upregulation in proliferating cells, and for their expression of an effector signature by upregulating expression of the key TFs *Prdm1*/BLIMP-1 on day 3 (our data and (42)), as well as E*pas1*/HIF-2a (57) and *Pparg* on day 5 (56,58). Early IL-4-dependent signaling was also essential to prevent IL-4Rα downregulation on activated CD4^+^ T cells (39), and thereby enabled TH2 cells to continue expanding according to the level of IL-4Rα-dependent signaling available and the competition for cytokine access, as also described for IL-2 (29). IL-4-dependent proliferation presumably required Ag^+^ APCs to support TCR engagement and cognate interactions, and was conceivably mediated through the induction of the transcriptional repressor Growth factor independent (GFI)-1 which supports proliferation and prevents apoptosis(10). IL-4-dependent proliferation may also contribute to stabilizing high GATA3 expression and the TH2 effector phenotype (73,74). Thus, while IL-4 may not direct TH2 differentiation, it powerfully regulates the magnitude of the TH2 response and TH2 progression towards effector function.

Our experiments did not directly examine the sources of the IL-2 and IL-4 that drive TH2 differentiation in our model, however, scRNAseq data provided some relevant information. On day 3, *Il2* was expressed by cells in several T cell clusters including cycling and memory cells, and was highly expressed by a population of CD69^+^Nr4a1^+^Egr3^+^ “Recently activated” CD4^+^ T cells, which were likely engaging in cognate interaction with Ag-presenting DCs and undergoing functional differentiation(75,76). A similar expression of CD69 and *Nr4a1*/NUR77 was also reported in OVA-specific OTII cells 24h after OVA immunization in TH2 conditions(33), whereas IRF4 expression was necessary for dense, LFA1 and ICAM1-dependent CD4^+^ T cell clustering at the T-B border to sustain TH2 differentiation(30). Strikingly, we found CD69^+^Nr4a1^+^Egr3^+^ cells in LN also on day 5, to suggest ongoing cognate T cell-APC interactions, but at this time they expressed *Il2* only if IL-4Rα^KO^, and *Il4* if IL-4Rα^WT^ to suggest an IL-4-dependent suppression of IL-2 expression as described (41,42,44). A CD69^+^Nr4a1^+^Egr3^+^ population was also found in *Ms*-immunized mice, but it was less abundant than following *Nb*, and was *Irf4*-low and *Il2*-low on day 3 possibly reflecting the need for TH1 cells to interact with APCs other than DCs in order to acquire effector function(77,78). In addition to T cells, CD301b^+^ DC2s also express functionally relevant *Il2* and modulate its bioavailability to CD4^+^ T cells through IL-2Rα expression(33).

*Il4* was expressed in an IL-4-independent manner by cycling cells and the WT and KO TFH clusters on day 3, while expression on day 5 was restricted to the WT TFH population to confirm that IL-4-dependent signaling was not necessary for initial *Il4*, but for sustained expression at later time points (79). This timing matched IL-4 protein expression as shown by the increased expression of IL-4Rα in bystander naïve cells in LN, which was already detectable on day 3 and was further upregulated on day 5, as well as by the IL-4-dependent expansion of the GATA3^hi^ TH2 population between day 3 and day 5.

In conclusion, our work proposes an alternative role for IL-4 in TH2 differentiation by showing that IL-4 and IL-4Rα expression were not necessary for initial GATA3 upregulation or the acquisition of TH2 identity, but were essential for TH2 proliferation in LN and to enable an effector TH2 program through the expression of TFs including BLIMP-1, EPAS1/HIF-2a and PPARG (35,56,70,80). Early IL-4 was also essential to sustain IL-4Rα expression in CD4^+^ T cells undergoing activation, thus maintaining high IL-4 cytokine responsiveness as well as the developing TH2’s ability to compete for IL-4 with bystander cells. Understanding how IL-4 access is regulated within tissue contexts will be essential to understanding its biological functions and impact on TH2 differentiation *in vivo*.

## Author Contributions

GRW and FR conceived and designed experiments; GRW, MSF, SCT, JC and EH carried out experiments; GRW, EH, MSF and FR analysed data; OL designed and interpreted scRNAseq and bioinformatics analyses; CC and SIO analyzed scRNAseq data, performed bioinformatic analyses and data visualization; JC and KLH contributed gene signatures. GRW, CC, OL and FR wrote the manuscript; all authors read and provided feedback on manuscript and figures.

## Acknowledgements

The Authors gratefully thank Samantha Small and Alix Grooby for performing cell sorting; Kate Pilkington for inexhaustible expertise and advice on spectral flow analysis; Sventja von Daake for scRNAseq library preparation; David Eccles and Cynthia Morgan for processing of FastQ files on the SevenBridges platform; all staff of the Hugh Green Technology Center for support; and all BRU staff for excellent animal care and husbandry. We are also grateful to Professor Frank Brombacher, Cape Town, for generously donating IL-4Rα^KO^ breeding pairs, and Professors Alex Dent, Indianapolis and Carola Vinuesa, Canberra, for CD4-Cre, Bcl6^fl/fl^ mice, and Professor Graham Le Gros of the Malaghan Institute for discussion.

This work was funded by an Independent Research Organisation grant from the Health Research Council (HRC) of New Zealand to the Malaghan Institute of Medical Research, and HRC project grants 18/510 and 21/186 to FR. GRW was funded by a PhD scholarship from the University of Otago Wellington, New Zealand.

## Materials and Methods

### Mice

All mice were bred and housed at the Malaghan Institute of Medical Research animal facility under specific pathogen-free conditions and were age- and sex-matched within experiments. C57BL/6J (C57, strain #000664) and B6.SJL-Ptprc^a^ Pepc^b^/BoyJ (B6.SJL, strain #002014) were originally obtained from the Jackson Laboratory (Bar Harbor, ME). Bcl6^fl/fl^.CD4-cre (BCL6^ΔCD4^) mice (81,82) were provided by the Australian Phenomics Facility, ANU, Canberra, Australia. IL-4Rα^fl/fl^ (83) and IL-4Rα^KO^ (84) mice on a C57BL/6 background were obtained from Prof Frank Brombacher, International Centre for Genetic Engineering and Biotechnology, Cape Town, South Africa. Experimental protocols were approved by the Victoria University of Wellington Animal Ethics Committee and were performed according to institutional guidelines.

### Bone Marrow chimeras

Recipient C57BL/6 x B6.SJL mice aged 10 –12 weeks were treated with two doses of 5.5 Gy using a Gammacell 3000 Elan irradiator (MDS Nordion) given 3h apart. One day after irradiation, mice received 5 – 10 x 10^6^ BM cells from sex-matched donor mice prepared by flushing donor femurs and tibias with sterile IMDM (Gibco) and filtering through a sterile 70 µm cell strainer (Falcon), as detailed in text and figure legends. Chimeric mice were housed in individually ventilated cages and supplied with 2 mg/mL Neomycin trisulfate (Sigma) -supplemented drinking water for the first 3 weeks. Chimeras were rested for at least 10 weeks before immunization.

### Adoptive transfer experiments

CD4^+^ T cells were purified from spleens of B6.SJL donor mice using the Dynabead® FlowComp™ Mouse CD4^+^ Enrichment kit (Invitrogen) and labelled at 1×10^6^ cells/mL with 3μM CellTrace Violet (ThermoFisher) in sterile PBS for 30 minutes at 37°C. A total of 3 – 5 x 10^6^ labelled cells were injected intravenously into C57 or IL-4Rα^fl/-^recipients as indicated. *In vivo* proliferation was assessed on day 5 after immunization.

### Immunizations and *in vivo* treatments

*Nippostrongylus brasiliensis* infective L3 larvae (*Nb*) were prepared, washed in sterile PBS and inactivated by three freeze-thaw cycles as previously described (85,86). *Mycobacterium smegmatis* (*Ms*; mc2155) was grown in LB broth (Difco LB Lennox – low salt, BD) at 37 °C overnight. Bacteria were washed three times in PBS 0.05% Tween 80, heat-killed at 75 °C for 1 h and stored at −70 °C. For immunizations, mice were anesthetized and injected with 300 *Nb*, 4×10^6^ *Ms* or 20μg HDM extract (Greer Laboratories, Lenoir, NC, USA) intradermally (i.d.) into the ear pinna as described (22). To block IL-4 signaling *in vivo,* mice were treated with 500µg αIL-4 (clone 11B11, rat IgG1, k) or mouse IgG1 (clone HRPN, anti-horseradish peroxidase rat IgG1, k) *InVivo*MAb (BioXCell) given i.p. following immunization as specified. To block IL-2 signaling *in vivo,* mice were treated with 500µg αIL-2 (clone JES6-1A12, rat IgG2a, k) or rat IgG2a (clone 2A3, anti-trinitrophenol rat IgG2a, k) *InVivo*MAb (BioXCell) given i.p. following immunization as specified.

### Cell Preparations

Lymph node and spleen single-cell suspensions were prepared by gently pressing through a 70µm cell strainer using the plunger of a 1mL syringe and rinsing with non-supplemented IMDM (Invitrogen). Red blood cells in spleen preparations were lysed using sterile RBC lysis buffer (Sigma). For assessment of T cell intracellular cytokines, LN single cell suspensions were cultured in foetal calf serum (FCS)-supplemented IMDM (Invitrogen) in the presence of 50ng/mL PMA (Sigma-Aldrich), 1µg/mL ionomycin (Merck Millipore) and 1µL/mL GolgiStop™ (BD Pharmingen) for 5 hours at 37°C.

### CRISPR/Cas9 gene editing

CD4^+^ T cells were purified from spleens of B6.SJL donor mice using the EasySep™ Mouse CD4^+^ T Cell Isolation Kit (STEMCELL) and electroporated using the Lonza 4D-NucleofectorTM, X Unit (Lonza, Cat# AAF-1002B) as previously described (87). Gene Knockout Kit v2 single-guide (sg)RNAs targeting the murine *Il2ra* gene (5’-UGGCAUUGGGGACCUCGGGU -3’; 5’- CUUGCAUUCACAGUUUAGGA-3’; 5’-UGCGUUGCUUAGGAAACUCC-3’), non-targeting scramble sgRNA (5’-GCACUACCAGAGCUAACUCA -3’) and spCas9 2NLS Nuclease were all from Synthego. Briefly, 5×10^6^ purified CD4^+^ T cells in 20µl P3 buffer (Lonza 4D Nucleofector kit) were mixed with sgRNA/Cas9 RNP complexes containing 300pmol sgRNA and 37.5pmol SpCas9 in a total volume of 5µl. The cell/RNP mix was transferred to the bottom of a Nucleocuvette^®^ Strip well and pulsed using the ‘mouse T cell unstimulated’ program (DN100). After 10min of recovery in complete medium, edited cells were washed, resuspended in 150ul of sterile PBS and injected intravenously into host mice as indicated. Successful editing was assessed by flow cytometry in each experiment.

### Flow Cytometry

Cell suspensions were prepared as described and stained in 96-well round-bottom plates (Falcon) with Zombie NIR (Biolegend) in PBS to identify non-viable cells. For staining of cell surface markers, cells were suspended in anti-mouse Fc receptors block (clone 2.4G2, affinity purified from hybridoma culture supernatant) prior to labelling with cocktails of fluorescent antibodies made up in PBS containing 2mM EDTA, 0.01% sodium azide and 2% FCS, supplemented with Brilliant Staining Buffer (BD). Chemokine receptor (CCR4, CXCR5…) staining was by incubation with the appropriate antibody concentrations for 30 minutes at 37°C prior to staining for cell surface antigens. Intracellular and intranuclear staining was performed using a FOXP3 Transcription Factor staining kit (eBioscience) following extracellular markers staining and three wash steps by addition of 200uL PBS supplemented with 2mM EDTA, 0.01% sodium azide and 2% FCS to each well, and centrifugation at 300g. Antibodies and streptavidin conjugates used in experiments are detailed in **Table 1**. Cell suspensions were washed three times before analysis; spectral reference controls were recorded using single cell suspensions from relevant tissues, or using UltraComp eBeads™ (eBioscience). Dead cells and doublets were excluded from analysis as shown in **Supplemental Figure 1**. Samples were collected on a 3- or 5- laser Aurora Spectral analyser (Cytek, Fremont, CA) and analysed using FlowJo software (version 10, Becton Dickinson, San Jose, CA).

**Table 1:** Antibodies and Streptavidin Conjugates used in this work.

| Specificity | Clone | Fluorochrome | Company | RRID |
| --- | --- | --- | --- | --- |
| **CD3** | 145-2C11 | BUV395 | BD | AB_2738278 |
| **CD4** | GK1.5 | BUV805 | BD | AB_2739008 |
|  |  | BV750 | Biolegend | AB_2734150 |
|  |  | APC-eFluor 780 | ThermoFisher | AB_11218896 |
|  | RM4-5 | BV605 | BD | AB_2687549 |
| **CD8** | 53-6.7 | APC-H7 | BD | AB_1645237 |
|  |  | Pacific Blue | BD | AB_397029 |
|  |  | Biotin | BD | AB_394773 |
|  | HL3 | Biotin | BD | AB_395059 |
| **CD19** | 6D5 | Alexa Fluor 488 | Biolegend | AB_493339 |
| **IL-2RA (IL-2Rα)** | PC61 | BV480 | BD | AB_2739593 |
|  |  | APC | BD | AB_398623 |
| **CD44** | IM7 | Alexa Fluor 700 | eBioScience | AB_494011 |
|  |  | BUV737 | BD | AB_2870126 |
| **CD45.1** | A20 | APC-eFluor 780 | eBioScience | AB_1582228 |
|  |  | BV510 | Biolegend | AB_2563378 |
|  |  | PE | eBioscience | AB_465675 |
| **CD45.2** | 104 | APC-Cy5.5 | Southern Biotech | AB_2795317 |
|  |  | FITC | BD | AB_395041 |
| **CD45R (B220)** | RA3-6B2 | Biotin | eBioScience | AB_466449 |
| **CD124 (IL-4Rα)** | mIL4R-M1 | Alexa Fluor 647 | BD | AB_2715465 |
| **CD185 (CXCR5)** | 2G8 | PerCP-eFluor710 | BD | AB_2573837 |
|  | SPRCL5 | Biotin | BD | AB_394301 |
| **CD194 (CCR4)** | 2G12 | PE-Cy7 | Biolegend | AB_2244410 |
| **CD279 (PD-1)** | 29F.1A12 | BV785 | Biolegend | AB_2563680 |
| **FcγR block** | 24G2 | Unconjugated | Produced in-house |  |
| **FOXP3** | FJK-16S | Alexa Fluor 532 | ThermoFisher | AB_11219265 |
| **GATA3** | L50-823 | BV711 | BD | AB_2739242 |
|  |  | Alexa Fluor 647 | BD | AB_1645316 |
| **IFNγ** | XMG1.2 | FITC | BD | AB_395375 |
| **IL-2** | JES6-5H4 | BV711 | Biolegend | AB_2564225 |
| **IL-4** | 11B11 | APC | BD | AB_395391 |
|  |  | PE | BD | AB_398556 |
|  |  | PE-Cy7 | Biolegend | AB_10898116 |
| **Ly6c** | HK1.4 | Biotin | Biolegend | AB_1236553 |
| **Ly6G** | 1A8 | Biotin | Biolegend | AB_1186105 |
| **pSTAT6**<br>**(Tyr641)** | 46H1L12 | N/A | ThermoFisher | AB_2532305 |
| **Rabbit IgG** | Polyclonal | Alexa Fluor<br>647 | ThermoFisher | AB_2535812 |
| **Streptavidin** | N/A | PE-CF594 | BD | AB_11154598 |
|  |  | BV421 | Biolegend |  |
| **Tbet** | 4B10 | PE | Biolegend | AB_2200542 |
| **TcRβ** | H57-597 | BUV395 | BD | AB_2740818 |
|  |  | BV605 | BD | AB_2687544 |
| **TcR$\gamma\delta$** | GL3 | APC | Biolegend | AB_2563356 |

### Cell sorting

Sterile single-cell suspensions were prepared from skin-draining LNs of mixed (IL-4Rα^WT^ + IL-4Rα^KO^) BM chimeras on either day 3 or 5 after immunization with *Nb* or *Ms,* as detailed above. Cells were incubated in 5mL round-bottom tubes (Falcon) with Zombie NIR in PBS for 25 minutes on ice, followed by 2.4G2 Fc block for 10 minutes on ice in PBS supplemented with 5% FCS. Cells were spun down and incubated with a sterile sorting antibody master mix (TcRb-BV605, CD3-BUV395, CD4-BV750, CD45.1-PE, CD45.2-FITC, CD44-AF700, PD-1-BV785, CD62L-PE-Cy7, TCRγδ-APC, and a biotinylated dump channel of B220, Ly6c, NK1.1CD8, CD11b, and CD11c with SAV-BV421) for 15 minutes on ice. For each immunization and time point, cells were sorted into IL4Ra^KO^ activated, IL4Ra^KO^ naïve, IL4Ra^WT^ activated and IL4Ra^WT^ naïve using a Cytek Aurora CS and the gating strategy in **Figure S1B**, for a total of 16 samples. Each sample included LN cells from three donor mice combined together in equal ratios. Sorter set-up and QC were performed on the day of the sort with single-stained cells or beads for spectral unmixing. 8000 activated and 2000 naive cells of each genotype were sorted into sterile 2mL lo-bind tubes (Eppendorf) that had been pre-coated with FCS and contained BD Pharmingen staining buffer supplemented with 5% FCS. Sorted populations were kept on ice prior to BD Rhapsody analysis.

### Bulk RNA sequencing of *Il4-*AmCyan+ CD4⁺ T cells from lymph nodes

*Il4-*AmCyan and *Il13*-dsRed (4C13R) dual reporter mice (62) were immunized intradermally in the ear pinna with 30 µL of a solution containing 200 µg of house dust mite (HDM) whole bodies (Greer Laboratories, Lenoir, NC, USA). Mice were sacrificed on day 7 post-immunization, and CD4⁺ T cell subsets were isolated from ear-draining lymph nodes by fluorescence-activated cell sorting using a BD Influx cell sorter. *Il4-*AmCyan⁺ TFH cells were defined as CD45⁺CD3^int^CD4⁺CD8⁻PD-1⁺CXCR5⁺*Il4*-AmCyan⁺, whereas *Il4-*AmCyan⁺ non-TFH cells were defined as CD45⁺CD3^int^CD4⁺CD8⁻CXCR5⁻*Il4*-AmCyan⁺. A total of 1,000 cells per population were sorted directly into 5 µL of TCL lysis buffer (Qiagen, Hilden, Germany) supplemented with 1% 2-mercaptoethanol, in duplicate. Smart-seq2 libraries were prepared as previously described (51). Gene signatures for *Il4-*AmCyan⁺ TFH and *Il4-*AmCyan⁺ non-TFH cells were defined by differential gene expression analysis comparing *Il4-*AmCyan⁺ TFH and *Il4-*AmCyan⁺ non-TFH populations using DESeq2.

### Single-cell RNA sequencing

Two separate BD Rhapsody Whole Transcriptome Amplification (WTA) libraries were prepared for *Nb* and *Ms* immunized mice. Sorted cells were sample tagged using the BD Rhapsody Multiplexing kit and protocol (Single Cell Labelling with the BD Single -Cell Multiplexing Kit ID:210970 Rev 3.0) based on oligo tagged antibodies (anti-mouse CD45, Clone 30-F11, BD Biosciences). All eight samples were pooled at a ratio of 1:4 naïve cells to activated cells. 4.0 ×10^4^ cells from the pool were captured using a BD Rhapsody Single-Cell Analysis System (Doc ID: 210966 Rev.3.0). Sequencing libraries were prepared using the BD Rhapsody TM Library Reagent Kit Library preparation mRNA Whole Transcriptome Analysis (WTA), Sample Tag Library Preparation Protocol Doc ID:23-24119(01). The quality of the final libraries was assessed using a TapeStation 4150 (Agilent) with High Sensitivity D5000 ScreenTape and quantified using a Quantus Fluorometer (Promega). Sequencing was performed at the Ramaciotti Centre for Genomics (University of New South Wales, Sydney), where libraries were sequenced on a NovaSeq 6000 (S2 100 cycles) at a depth of ∼50,000 reads/cell for the WTA libraries and ∼600 reads/cell sample tags. FastQ files and project metadata were then processed and quality controlled using the BD Rhapsody WTA Analysis Pipeline (BD Biosciences) implemented on the SevenBridges platform. Reads were quality filtered, bead barcodes and UMIs were extracted and error-corrected, and sequences were aligned to the mouse reference genome GRCm39 with ENSEMBL gene annotation supplied in the BD Rhapsody Mouse WTA Reference Bundle. UMIs were collapsed to obtain raw gene-by-cell count matrices.

### Clustering and annotation

The Seurat V5 package (88) was used to process unfiltered single cell matrix, reduce dimensionality, cluster cells, and perform differential expression analysis. Multiplets, undetermined cells, cells with > 25% mitochondrial reads or < 300 feature counts were excluded. Doublets were identified and removed using the *DoubletFinder* package (89). Data were normalized, scaled, and variance-stabilized using SCTransform, regressing out cell cycle effects (90). Datasets from *Nb*-immunized and *Ms*-immunized samples were integrated using the Harmony package (91) to correct for batch effects. Dimensionality was reduced with PCA, and Harmony UMAP embeddings were generated for downstream analyses. Clustering was performed using the Louvain algorithm, with cell identities assigned on the basis of marker gene expression identified using the *FindAllMarkers* function (Wilcoxon rank-sum test with Benjamini–Hochberg correction). Cluster abundance differences were assessed using a permutation test implemented in the *scProportionTest* package, with significance defined at log2 fold change (log2FC) > 0.58 and false discovery rate (FDR) < 0.05. Differential gene expression (DEG) analysis was performed using the *FindMarkers* function (Wilcoxon rank-sum test with Benjamini–Hochberg correction) across T cell clusters. DEGs were identified based on an average log2FC > 0.58 and adjusted *P*-value < 0.05.

Gene expression values were additionally imputed using the Markov Affinity-based Graph Imputation of Cells (MAGIC) algorithm (92), a graph-based diffusion imputation approach to display smoothed expression distributions across cell populations, which was separately performed on IL-4Rα^WT^ or IL-4Rα^KO^ *Nb* datasets after normalization and dimensionality reduction.

### Gene and pathway enrichment analyses

Gene set enrichment was performed using single-sample gene set enrichment analysis (ssGSEA) as implemented in the *escape* package (v.2.2.2) (93). Differential enrichment analyses were performed separately depending on the gene set. For the stem-like and pathogenic TH2 signatures, enrichment between WT and IL-4Rα^KO^ cells was assessed using Wilcoxon rank-sum tests applied directly to the raw ssGSEA score matrix, with significance determined at FDR < 0.05. For the annotated TH2 and TFH signatures, enrichment was tested in a “one-vs-all” framework within *Nb*- or *Ms*-stimulated samples, highlighting whether a given cluster was significantly enriched for a signature compared to all other clusters. In this case, differential enrichment was defined as log2 fold-change ≥ 0.05 combined with FDR < 0.05, based on Wilcoxon rank-sum tests with Benjamini–Hochberg correction. Pathway enrichment analysis was performed on *Nb*- or *Ms*-stimulated Teff + Cycling-Teff or TFH-1 cells using the STRING database (94) against multiple databases, including KEGG, Gene Ontology (GO), and Reactome. Pathways with FDR < 0.05 were considered significantly enriched. For ranking, we applied the STRING *strength* parameter, defined as log10 (observed/expected). This metric quantifies the magnitude of enrichment by comparing the number of genes in the network annotated with a given term to the number expected in a random network of the same size.

### Transcription Factor (TF) regulatory network inference

TF regulatory network analysis was performed using pySCENIC (version 1.12.1) (95) on mouse scRNA-seq data from *Nb*- and *Ms*-immunized sample. Gene regulatory networks were inferred from the expression matrix using GRNBoost2 (*arboreto,* version 0.1.6) with a curated mouse TF list as candidate regulators. Co-expression modules were pruned to motif-supported regulons using cisTarget/ctxcore mm10 v10 ranking databases and motif annotations (motifs-v10nr; MGI), applying standard enrichment thresholds (NES-based pruning). Regulon activity was quantified per cell and per cluster using an AUC matrix generated using *AUCell*. Differential regulon activity between ^WT^ and KO cells (overall or within clusters/timepoints) was assessed using two-sided Mann–Whitney U tests with Benjamini–Hochberg FDR correction. Regulon activity was visualized using Z-scored cluster-level mean AUCell heatmaps.

### RNA-velocity, trajectory and pseudotime analysis

Spliced and unspliced read counts were generated with *velocyto.py* (96) using the mouse genome (GRCm39), generating loom files. RNA velocity analysis was performed on QC-filtered of WT and IL-4Rα^KO^ CD4⁺ T cells after *Nb* infection. After subsetting, data were re-processed to remove low-quality cells and re-normalized to retain highly variable genes with sufficient spliced/unspliced information for velocity modeling. RNA velocity was estimated using an expectation-maximization (EM) model implemented in the *scVelo* package (97). Velocity vectors were projected onto UMAP embeddings. Pseudotime analysis was performed in R studio using Monocle3 (v 1.3.7) (98). Treg cells were removed and analysis was restricted to the *Nb*-stimulated CD4⁺ T cell subset. A Monocle3 cell_data_set was generated from the Seurat RNA raw UMI count matrix and associated metadata. Data were pre-processed and clustered using the pre-existing UMAP embedding, and ordered in pseudotime using the Naïve CD4 T-cell population as the initial root. Pseudotime was computed on the combined WT and IL-4Rα^KO^ dataset. Genotype differences in pseudotime were assessed using two-sided Wilcoxon rank-sum tests, with effect sizes quantified using Cliff’s δ (99), interpreted according to standard magnitude categories (negligible, small, medium, large). Cluster-wise comparisons were corrected using the Benjamini–Hochberg method.

### Use of Large Language Model

Large Language Models (ChatGPT, versions 4o and 5; OpenAI) were used to assist with code generation, annotation and troubleshooting of R- and Python-based tools. All outputs were reviewed, tested, and validated by the authors.

### Statistical Analyses

Statistical analyses were performed using Prism 10 GraphPad software. For normally distributed data, comparisons between treatment groups were by ordinary two-way ANOVA with full model effects and Tukey’s multiple comparisons test, or as indicated in figure legends. Comparisons between two groups were with a Student’s *t*-test or as indicated. *P* values lower than 0.05 were considered significant and are referred to as follows: **** = *P* < 0.0001; *** = 0.0001 ≤ *P* < 0.001; ** = 0.001 ≤ *P* < 0.01; * = 0.01 ≤ *P* < 0.05; ns (not significant) = *P* ≥ 0.05.

