## Supplemental Figures 1-8 for "IL-2- and IL-4-dependent signaling play separate and sequential roles in the differentiation of GATA3^hi^ TH2 cells *in vivo*"

**A: Gating for Activated, Naïve (PD-1$^{low}$, CD44$^{low}$), T-bet+, TFH and GATA3-hi cells**

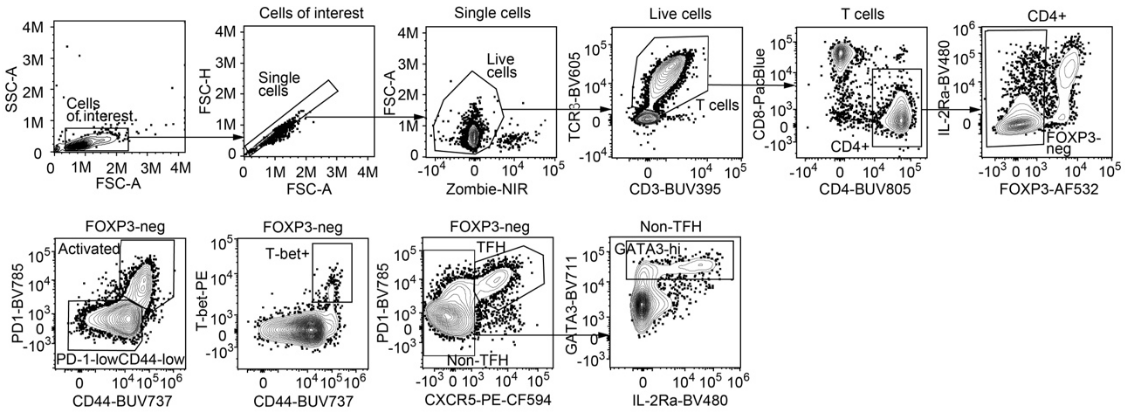


**B: CD4+ effector gating for cell division experiments**

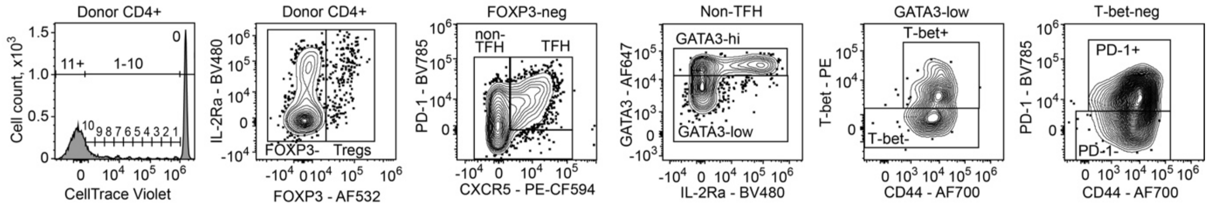


**C: Sorting strategy for scRNAseq**

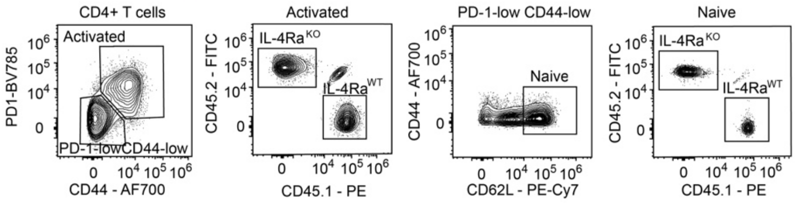


**D: Adoptive transfer experiments**

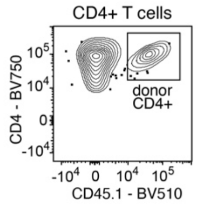


**Supplementary Figure 1: Gating strategies for CD4$^{+}$ T cell populations from immunized mice as indicated.**

Additional gatings are shown in the relevant figures.

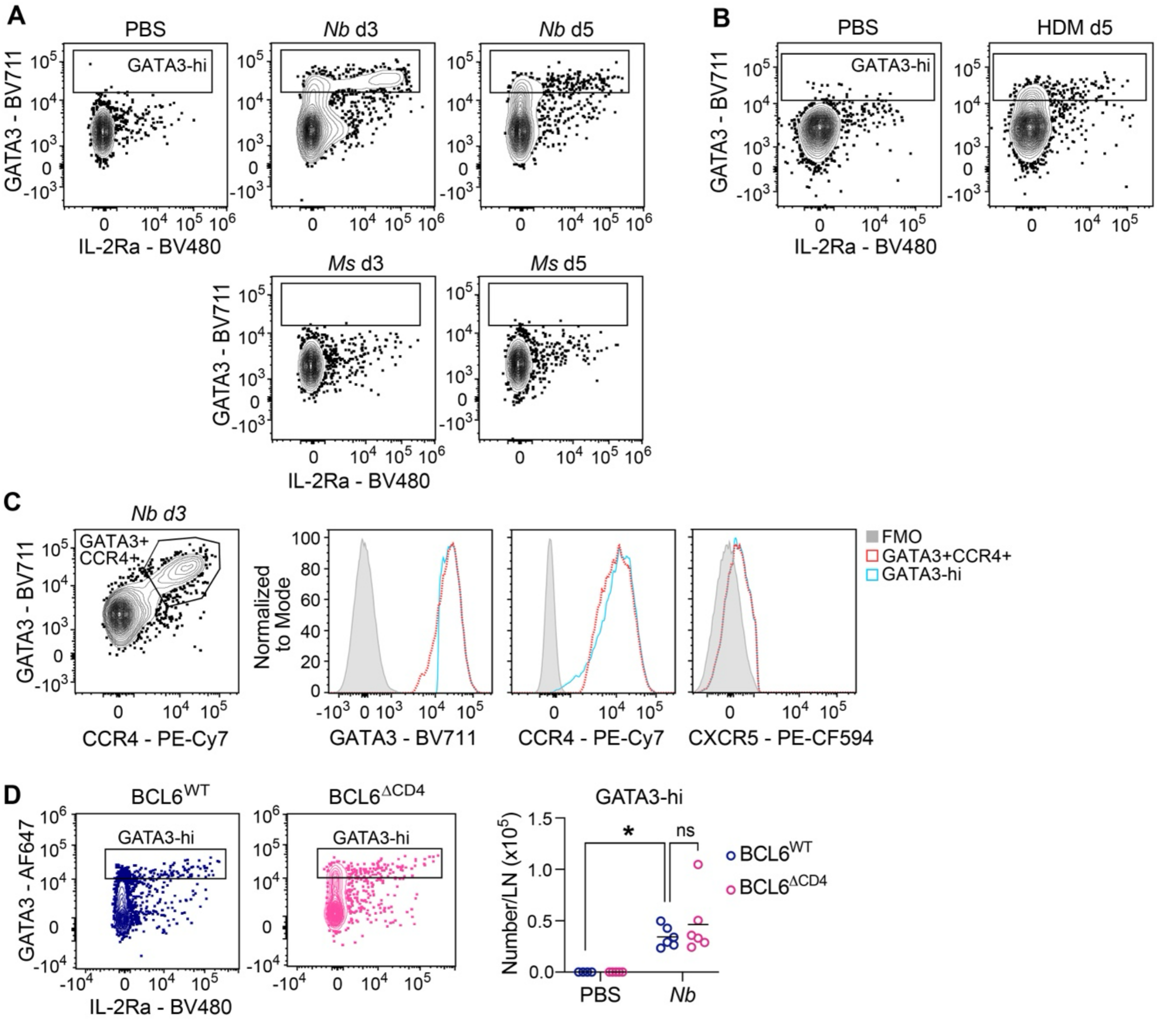


**Supplementary Figure 2: Intracellular transcription factor staining identifies GATA3$^{hi}$ TH2 cells in *Nb*- and HDM-immunized mice.**

C57BL/6 mice were immunized intradermally with *Nb*, *Ms* or House Dust Mite (HDM) and CD4$^{+}$ T cell responses were examined by intranuclear staining and flow cytometry at the indicated times after immunization.

**A:** GATA3$^{hi}$ (high) cells are found in lymph node following *Nb* or (**B**) HDM immunization, but not after PBS or *Ms*.

**C**: GATA3$^{hi}$ cells are CCR4$^{+}$ and do not express CXCR5. FMO: Fluorescence-minus-one.

**D:** GATA3$^{hi}$ cells are found in BCL6 conditional-KO mice that do not express BCL6 in CD4$^{+}$ T cells.

Panels show data from non-TFH cells gated as in Fig. S1A; *P* values in D were calculated by 2-way ANOVA with Tukey's correction; *: $P<0.05$; ns: $P>0.05$. Data are combined from two repeat experiments each with 3 replicates per group.

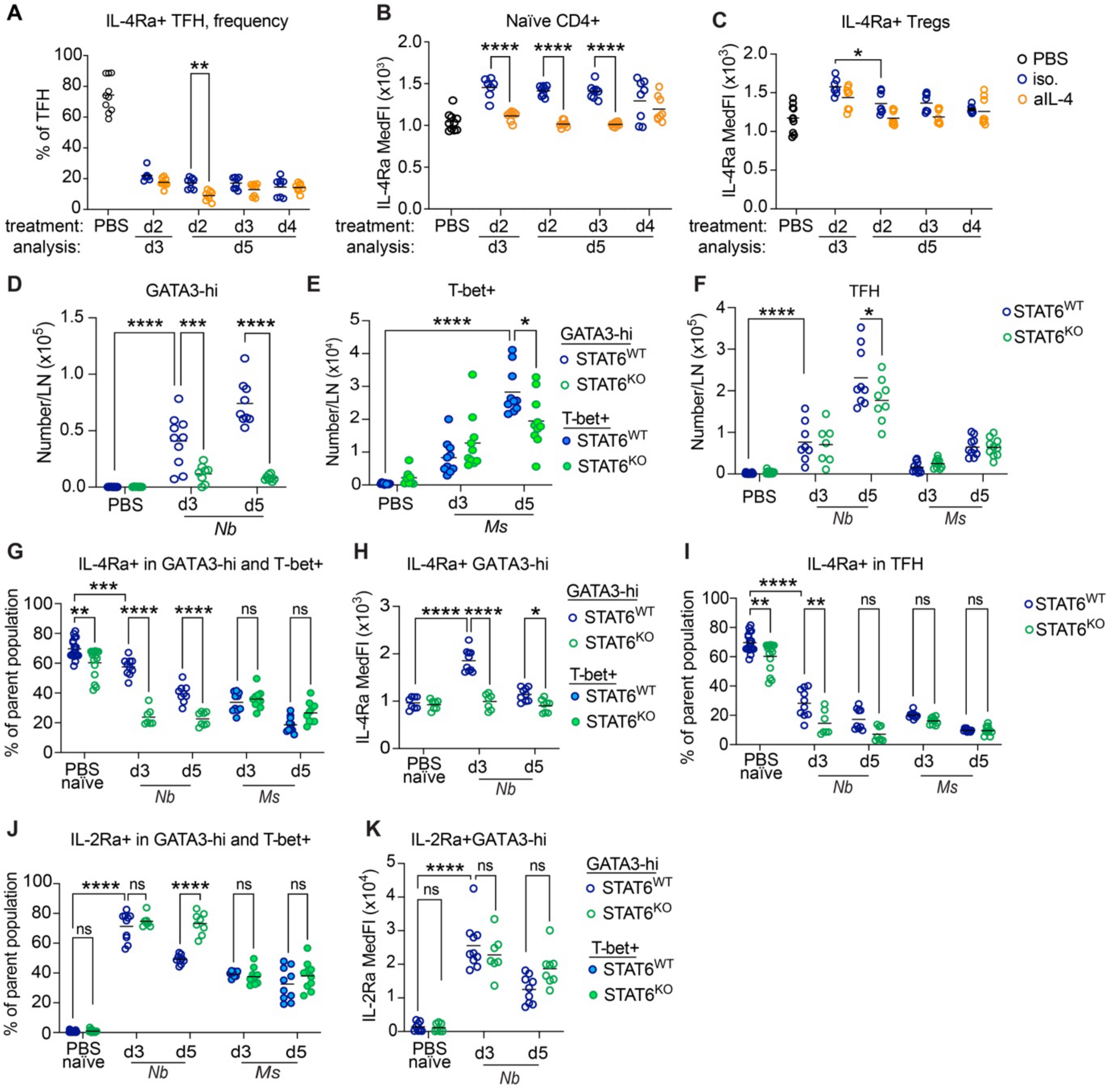


**Supplementary Figure 3: IL-4Rα and IL-2Rα expression on $CD4^+$ T cell subsets from *Nb*- or *Ms*-immunized mice.**

C57BL/6 mice treated with anti-IL-4 (aIL-4) or isotype control as in Figure 1A (**A-C**) and $STAT6^{WT}$ (C57BL/6) and $STAT6^{KO}$ mice (**D-K**) were immunized with *Nb* or *Ms*. $CD4^+$ T cell responses in the draining LN were examined by flow cytometry on day 3 and 5 after immunization as indicated.

**A:** Frequencies of IL-$4R\alpha^+$ cells in TFH populations from immunized and aIL-4-treated C57BL/6 mice.

**B:** IL-4Rα Median fluorescence intensities (MedFI) in naïve $CD4^+$ and (**C**) $FOXP3^+$ Treg cells from immunized and aIL-4-treated C57BL/6 mice.

**D:** Numbers of $GATA3^{hi}$ (**E**) T-$bet^+$ and (**F**) TFH cells per LN in *Nb*- or *Ms*- immunized $STAT6^{WT}$ and $STAT6^{KO}$ mice.

**G:** Frequencies of IL-$4R\alpha^+$ cells, and (**H**) MedFI of their IL-4Rα expression in the $GATA3^{hi}$ and $Tbet^+$ populations from immunized $STAT6^{WT}$ and $STAT6^{KO}$ mice.

**I:** Frequencies of IL-$4R\alpha^+$ TFH in $STAT6^{WT}$ and $STAT6^{KO}$ immunized mice.

**J:** Frequencies of IL-2Rα$^{+}$ cells, and (**K**) MedFI of their IL-2Rα expression in GATA3$^{hi}$ and Tbet$^{+}$ CD4$^{+}$ T cell populations from STAT6$^{WT}$ and STAT6$^{KO}$ immunized mice.
All data are pooled from two independent experiments, each with 3 – 5 mice per group, which gave similar results. *P* values were determine using two-way ANOVA with Tukey's multiple comparisons test. Each symbol refers to one mouse.
****, $P<0.0001$; ***, $P<0.001$; **, $P<0.01$; *, $P<0.05$; ~~ns, $P\geq0.05$;~~ selected *P* values are shown.

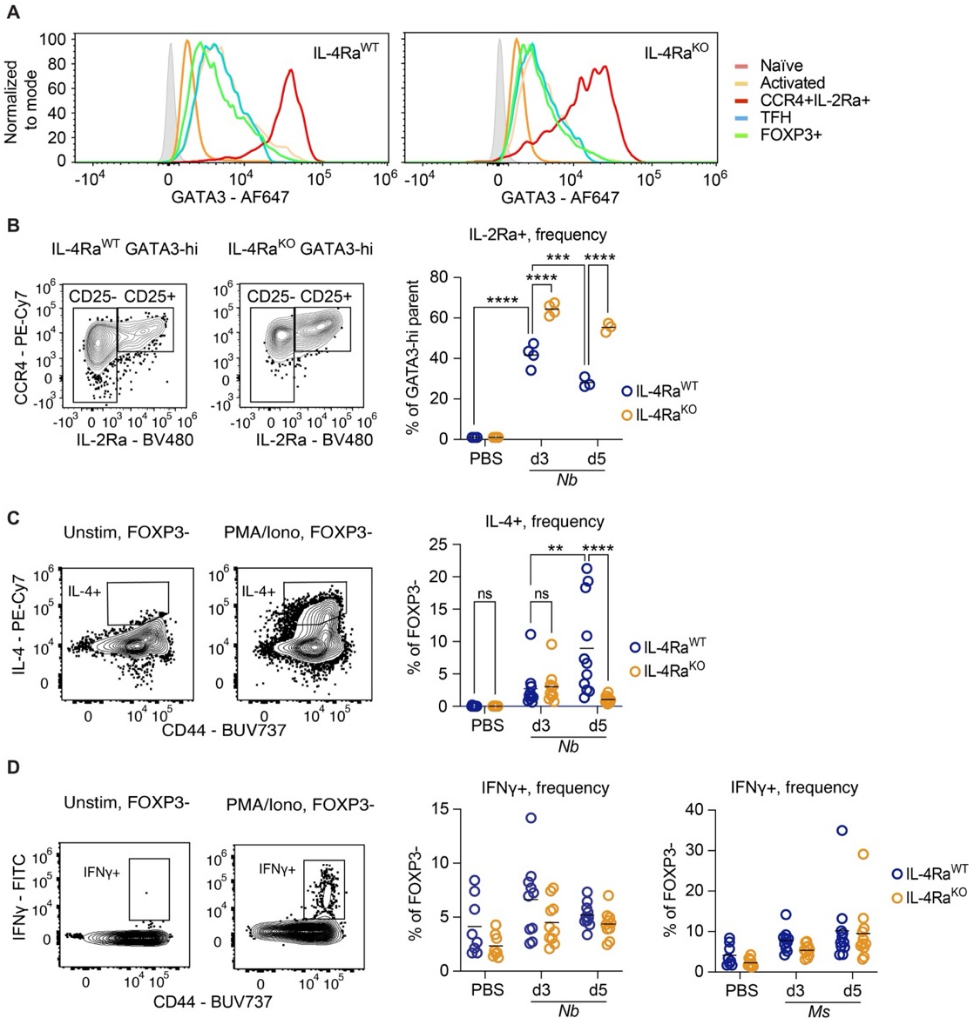


**Supplementary Figure 4: CD4$^+$ T cell phenotype and cytokine expression in (IL-4Rα$^{KO}$ + IL-4Rα$^{WT}$) mixed BM chimeras.**

Mixed (IL-4Rα$^{KO}$ + IL-4Rα$^{WT}$) BM chimeras set up as in Figure 2A were immunized intradermally with *Nb* or *Ms*. CD4$^+$ T cell responses in the draining LN were examined by flow cytometry on day 3 and 5 after immunization as indicated, using either surface and intranuclear staining (**A, B**) or intracellular cytokine staining following in vitro restimulation with PMA/ionomycin (**C, D**).

**A**: GATA3 expression in CD4$^+$ T cell populations from mixed BM chimeras on day 5 following ***Nb*** immunization.

**B:** Representative gating and frequencies of IL-2Rα$^+$ cells within the GATA3$^{hi}$ CD4$^+$ T cell population. Flow plots refer to day 5 following *Nb* immunization.

**C:** Representative gating and frequencies of IL-4$^+$ CD4$^+$ T cells. Flow plots refer to day 5 following *Nb* immunization.

**D:** Representative gating and frequencies of IFNγ$^+$ CD4$^+$ T cells. Flow plots refer to day 5 following *Nb* immunization.

Data refer to one representative of two (**A, B**) or two pooled (**C, D**) experiments, each with 3 - 5 mice per group, which gave similar results.

*P* values were calculated using two-way ANOVA with Tukey's multiple comparisons test. ****, *P*<0.0001; ***, *P*<0.001; **, *P*<0.01; *. Selected *P* values are shown.

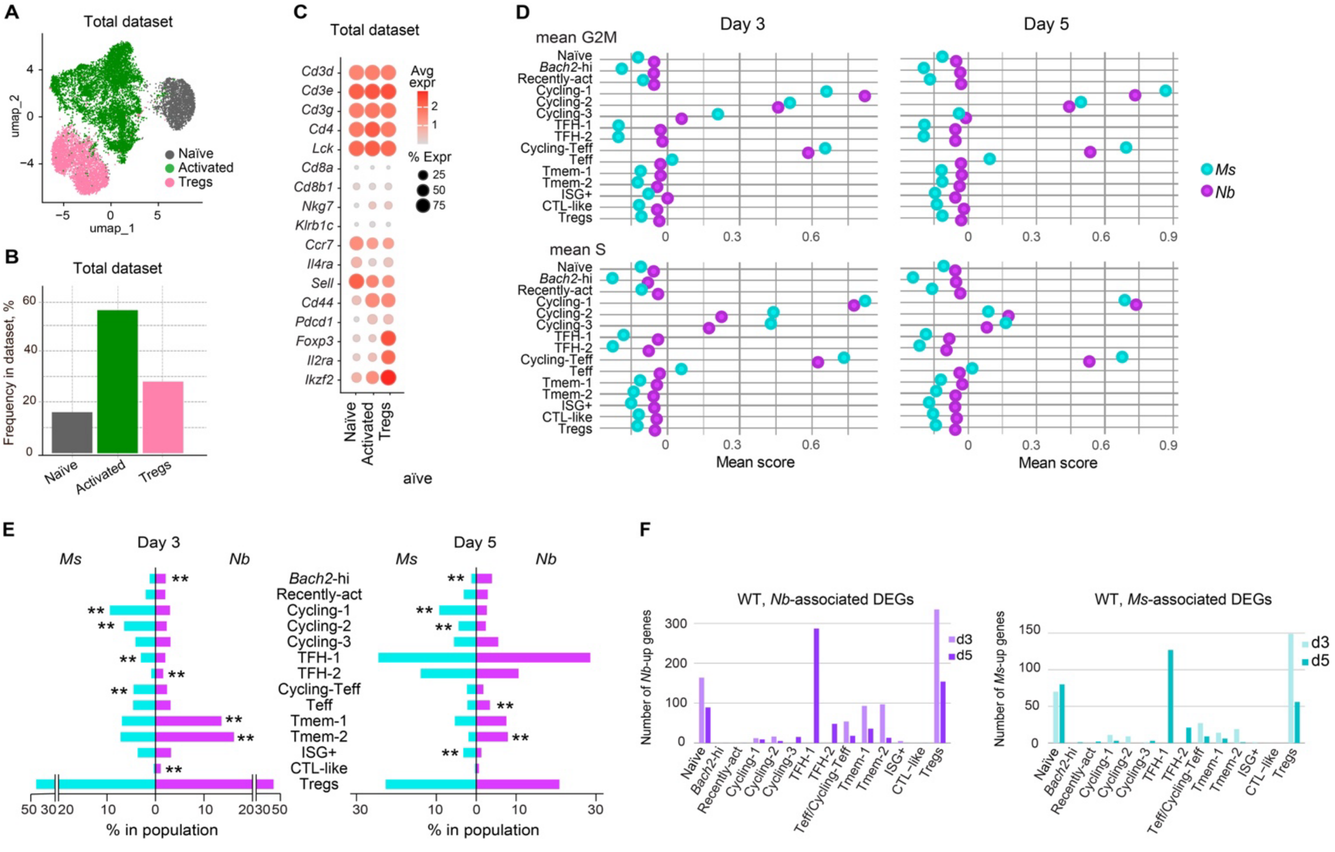


**Supplementary Figure 5: scRNAseq clustering and dataset characterization.**

Single-cell RNA sequencing of $CD4^+$ T cell populations from mixed (IL-4Rα$^{KO}$ + IL-4Rα$^{WT}$) BM chimeras on day 3 and 5 following *Nb* or *Ms* immunization.

**A:** Uniform manifold approximation and projection (UMAP) visualizing $CD4^+$ T cell populations in the dataset, and **B**: relative population abundance in the total (including *Nb* and *Ms*, day 3 and day 5, IL-4Rα$^{WT}$ and IL-4Rα$^{KO}$) dataset.

**C:** Expression of relevant transcripts in the $CD4^+$ T cell populations identified in **A**.

**D:** Mean G2M and S cell-cycle phase scores for the indicated clusters in the total dataset are shown according to immunization and time of analysis. Clusters were defined as in Fig. 3A.

**E:** Relative cluster abundance within the combined IL-4Rα$^{WT}$ and IL-4Rα$^{KO}$ "Activated" population in **A**; the Treg population is included as a reference.

Significance was determined by permutation testing: the observed difference between *Nb* and *Ms* was compared to the difference expected by chance, and *P* values were adjusted for multiple testing with the Benjamini–Hochberg FDR procedure. Significant values were defined as: **: FDR < 0.01.

**F:** Numbers of differentially expressed genes (DEGs) in IL-4Rα$^{WT}$ that were associated with *Nb* or *Ms* in an *Nb* vs *Ms* comparison, by cluster and day of immunization. DEGs were identified using a Wilcoxon rank-sum test with Benjamini–Hochberg correction, with significance defined as log2FC $\geq 0.585$ and FDR < 0.05.

scRNAseq data are from one (*Ms*) or two pooled (*Nb*) biological replicates.

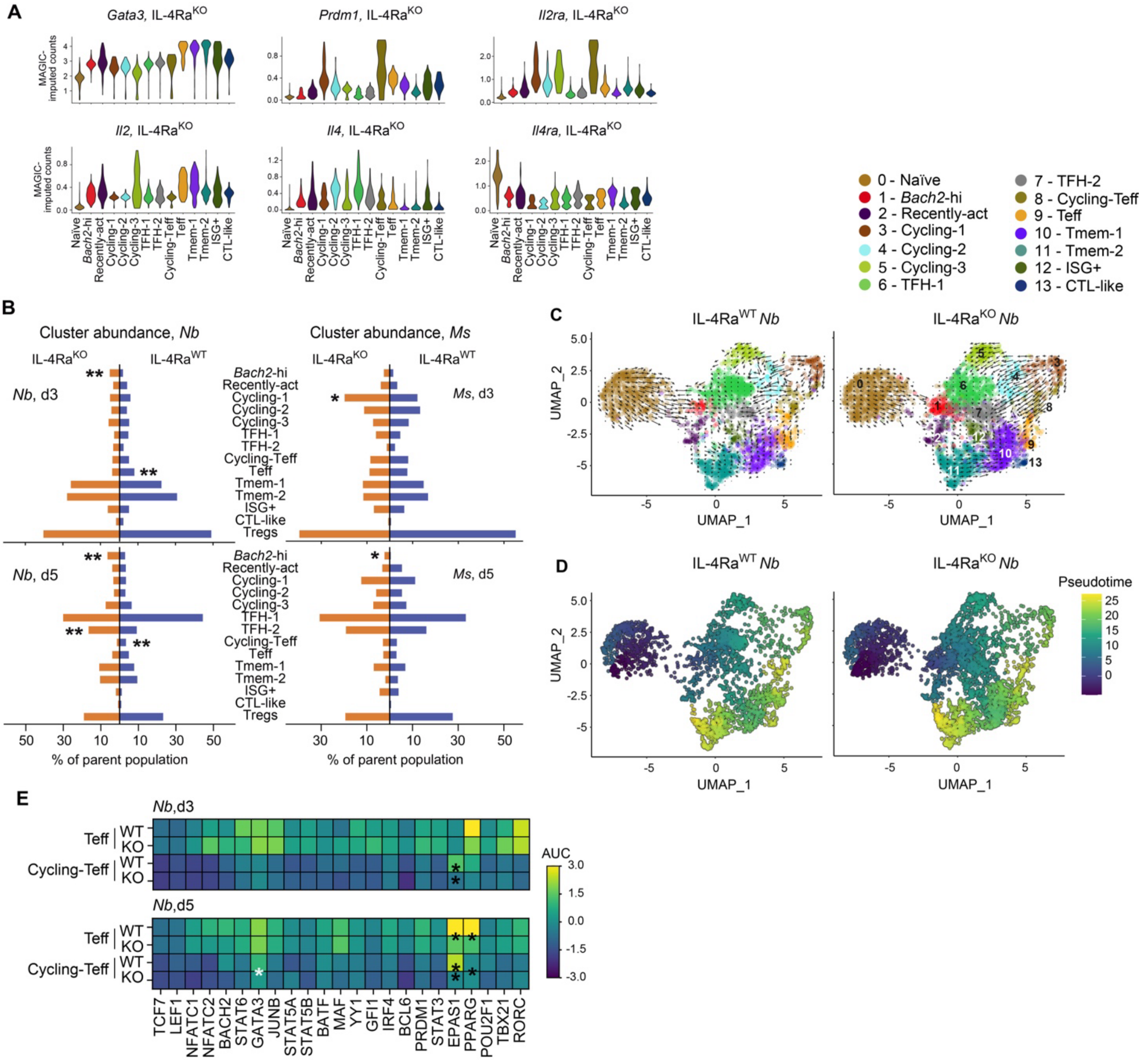


**Supplementary Figure 6: Comparative analysis of IL-4Rα$^{WT}$ and IL-4Rα$^{KO}$ populations in mixed BM chimeras following *Nb* or *Ms* immunization.**

Single-cell RNA sequencing analysis of CD4$^{+}$ T cell populations from (IL-4Rα$^{KO}$ + IL-4Rα$^{WT}$) mixed BM chimeras on day 3 and 5 following *Nb* or *Ms* immunization.

**A:** Violin plots showing single-cell expression levels of the indicated transcripts were generated using imputed expression values obtained with the MAGIC (Markov Affinity-based Graph Imputation of Cells) algorithm. Violins show combined day 3 and day 5 data in the IL-4Rα$^{KO}$ population from *Nb*-immunized chimeras.

**B:** Cluster abundance in IL-4Rα$^{KO}$ and IL-4Rα$^{WT}$ populations. Clusters were defined as in Figure 3A and significance was determined by permutation testing: the observed difference was compared to the difference expected by chance, and *P* values were adjusted for multiple testing with the Benjamini–Hochberg FDR procedure. Significant values were defined as log2FC > 0.58 and FDR < 0.05. **: FDR < 0.01; *: FDR < 0.05.

**C:** RNA velocity trajectories for IL-4Rα$^{WT}$ and IL-4Rα$^{KO}$ populations following immunization with *Nb* (day 3 and day 5 combined, excluding Tregs). RNA velocity vectors (black arrows) were computed from spliced, unspliced and ambiguous read counts with *velocyto.py*,

estimated using the scVelo expectation–maximization model. Vectors were projected onto the UMAP to visualize the inferred directionality of transcriptional state transitions across clusters.
**D:** Monocle3 pseudotime analysis of IL-4Rα$^{WT}$ and IL-4Rα$^{KO}$ populations (day 3 and day 5 combined, excluding Tregs) following *Nb* immunization. UMAP embeddings are shown colored by Monocle3 pseudotime using the Naïve CD4 T-cell population as the root.
**E:** Heatmap showing the regulon activity (z-scored area under the curve, AUC) for selected transcription factors in the indicated clusters**.** Differential regulon activity between IL-4Rα$^{WT}$ and IL-4Rα$^{KO}$ cells within each cluster was assessed using a two-sided Mann–Whitney U test with Benjamini–Hochberg FDR correction for multiple comparisons. Asterisks indicate statistical significance (**P-adj* < 0.05; ***P-adj* < 0.01, ****P-adj* < 0.001).
Data are from one (*Ms*) or two pooled (*Nb*) biological replicates.

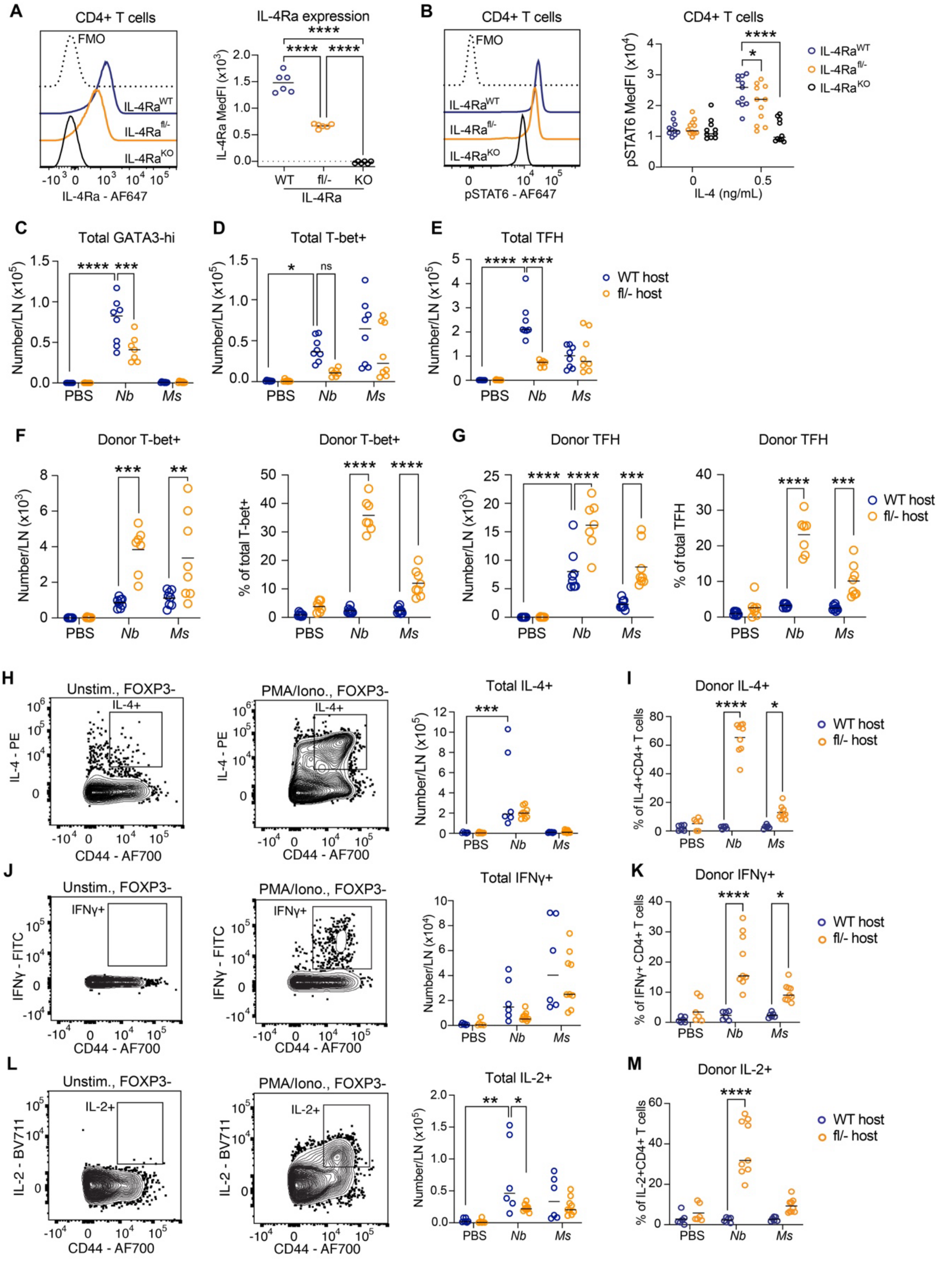


**Supplementary Figure 7: Reduced competition for IL-4 leads to magnified expansion of CD4$^+$ effector T cells in lymph node.**

**A, B:** Lymph node (LN) cells from the indicated mouse strains were examined for IL-4Rα expression and STAT6 phosphorylation by flow cytometry.

**A:** Representative histograms and Median fluorescence intensities (MedFI) of IL-4Rα expression on IL-4Rα$^{WT}$, IL-4Rα$^{fl/-}$ and IL-4Rα$^{KO}$ total CD4$^{+}$ T cells freshly isolated from mouse LN.
**B:** Representative histograms and MedFI of pSTAT6 in IL-4Rα$^{WT}$, IL-4Rα$^{fl/-}$ and IL-4Rα$^{KO}$ LN CD4$^{+}$ T cells cultured in medium supplemented with the indicated doses of IL-4.
**C - K:** IL-4Rα$^{WT}$ and IL-4Rα$^{fl/-}$ hosts were adoptively transferred with CellTrace Violet (CTV)-labelled IL-4Rα$^{WT}$ CD4$^{+}$ T cells and immunized with *Nb* or *Ms* as in the schematic in Figure 6A. CD4$^{+}$ T cell responses were measured by flow cytometry 5 days following immunization. Intracellular cytokine staining was performed after 5h *ex vivo* restimulation with PMA/ionomycin.
**C:** Total numbers (host + donor) of GATA3$^{hi}$; (**D**) T-bet$^{+}$ and (**E**) TFH cells in adoptively transferred mice.
**F:** Numbers and frequencies of donor T-bet$^{+}$ and (**G**) donor TFH in the respective total populations.
**H**: Representative gating and numbers of IL-4$^{+}$ T cells in adoptively transferred mice, and (**I**) frequencies of donor cells within the IL-4$^{+}$ population.
**J**: Representative gating and numbers of IFNg$^{+}$ T cells in adoptively transferred mice, and **(K)** frequencies of donor cells in the IFNg$^{+}$ CD4$^{+}$ T cell populations.
**L:** Representative gating and numbers of IL-2$^{+}$ T cells in adoptively transferred mice, and **(M)** frequencies of donor cells in the IL-2$^{+}$ CD4$^{+}$ T cell populations.
Data are pooled from two repeat experiments, each with 3 – 5 mice per group, which gave comparable results. Each symbol corresponds to one mouse. *P* values were calculated by two-way ANOVA with Tukey's multiple comparisons test. Selected significant P values are shown. ****, $P<0.0001$; ***, $P<0.001$; **, $P<0.01$; *, $P<0.05$.

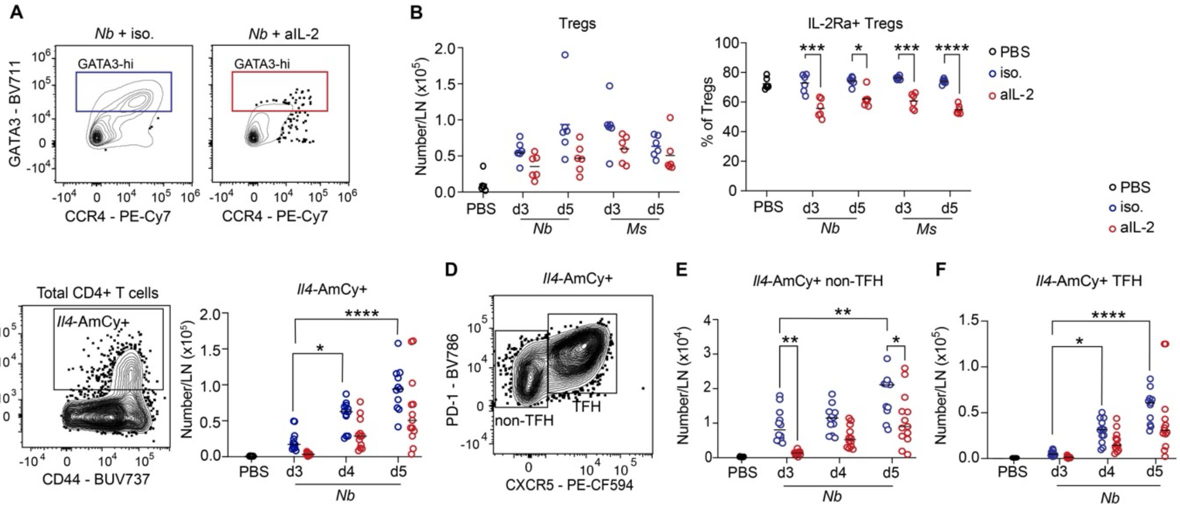


**Supplementary Figure 8: Characterization of $CD4^+$ T cell populations following aIL-2 treatment.**

C57BL/6 and 4C13R dual-reporter mice were immunized with *Nb* or *Ms*, and treated with neutralizing anti-IL-2 (aIL-2) antibodies or isotype control on either day 0 (**A-F**) or day 2 (**I+**) after immunization. $CD4^+$ T cell response in the draining LN were assessed by flow cytometry on day 3, 4 or 5 after immunization as indicated.

**A:** CCR4 and GATA3 expression in isotype or anti-IL-2-treated C57BL/6 mice on day 3 following *Nb* immunization.

**B:** Numbers of Tregs and frequencies of their IL-2Rα expression in immunized C57BL/6mice.

**C:** Representative gating and numbers of *Il4*-$AmCyan^+$ $CD4^+$ T cells in immunized 4C13R mice. Gating shows data on day 5.

**D:** Representative gating and numbers of (**E**) non-TFH and (**F**) TFH in the *Il4*-$AmC^+CD4^+$ T cell population in **B**. Data in **C** refer to day 5.

Data are pooled from 2 or 3 repeat experiments, each with 3 – 7 mice per group, which gave similar results.

*P* values refer to the comparison between treatment groups and were calculated using ordinary two-way ANOVA with Tukey's multiple comparisons test. Only selected *P* values are shown. ****, $P<0.0001$; ***, $P<0.001$; **, $P<0.01$; *, $P<0.05$.
